# Read2Tree: scalable and accurate phylogenetic trees from raw reads

**DOI:** 10.1101/2022.04.18.488678

**Authors:** David Dylus, Adrian Altenhoff, Sina Majidian, Fritz J Sedlazeck, Christophe Dessimoz

**Affiliations:** Department of Computational Biology, University of Lausanne, 1015 Lausanne. Switzerland; F. Hoffmann-La Roche Ltd, Immunology, Infectious Disease, and Ophthalmology (I2O), Roche Pharmaceutical Research and Early Development (pRED), Basel, 4070, Switzerland; SIB Swiss Institute of Bioinformatics, 1015 Lausanne, Switzerland; Department of Computer Science, ETH, 8092 Zurich, Switzerland; Human Genome Sequencing Center, Baylor College of Medicine, Houston, TX, 77030, USA; Department of Computer Science, Rice University, Houston, TX, 77005, USA; Department of Computer Science, University College London, London WC1E 6BT UK; Centre for Life’s Origins and Evolution, Department of Genetics, Evolution and Environment, University College London, London WC1E, UK

## Abstract

The inference of phylogenetic trees is foundational to biology. However, state-of-the-art phylogenomics requires running complex pipelines, at significant computational and labour costs, with additional constraints in sequencing coverage, assembly and annotation quality. To overcome these challenges, we present Read2Tree, which directly processes raw sequencing reads into groups of corresponding genes. In a benchmark encompassing a broad variety of datasets, our assembly-free approach was 10-100x faster than conventional approaches, and in most cases more accurate—the exception being when sequencing coverage was high and reference species very distant. To illustrate the broad applicability of the tool, we reconstructed a yeast tree of life of 435 species spanning 590 million years of evolution. Applied to *Coronaviridae* samples, Read2Tree accurately classified highly diverse animal samples and near-identical SARS-CoV-2 sequences on a single tree—thereby exhibiting remarkable breadth and depth. The speed, accuracy, and versatility of Read2Tree enables comparative genomics at scale.

## INTRODUCTION

Phylogenetic trees depict evolutionary relationships among biological entities. These entities can be species—as in the Tree of Life^1–4^. They can also be cancerous cells in tumour progression trees^5^ or developmental lineage trees^6^, viral and bacterial strains in infectious outbreaks^7^, cells, or genes in trees used to propagate molecular function annotations among model and non-model species^8,9^. Because of this pervasiveness, methods to infer phylogenetic trees are among the most used and cited software tools in all of life sciences.

In the context of species tree inference, the availability of genome-wide sequencing has made it routine to consider as many marker genes per taxon as the genomes provide. This “phylogenomic” approach has resolved many key aspects of the eukaryotic tree of life, such as the relation among deep angiosperm clades^10^, the position of sea squirts within chordates^11^, the Ecdysozoa clade^12^, the Lophotrochozoa clade^13^, relations among main myriapod clades^14^, among many others.

Nevertheless, despite rapid improvements in quality and cost of sequencing^15,16^, the data analysis required to infer phylogenetic trees remains extremely labour and computational intensive^17^. Phylogenomic studies require multiple costly steps, each of which can be major research endeavours (**Fig 1**): the curation of raw reads, the *de novo* assembly often including multiple rounds of error corrections and scaffolding either with one or multiple technologies^18^, the annotation and characterization of important genes, the identification and comparison of orthologous genes, and the tree inference from orthologous markers. The current best practices optimise this process using costly technology combinations—such as long and short read sequencing—and multiple rounds of parameter optimizations across multiple pipelines. Still, the problem remains compute intensive and requires different skill sets from different areas of specialisation (e.g. assembly, annotation, phylogeny).

**Figure 1.**
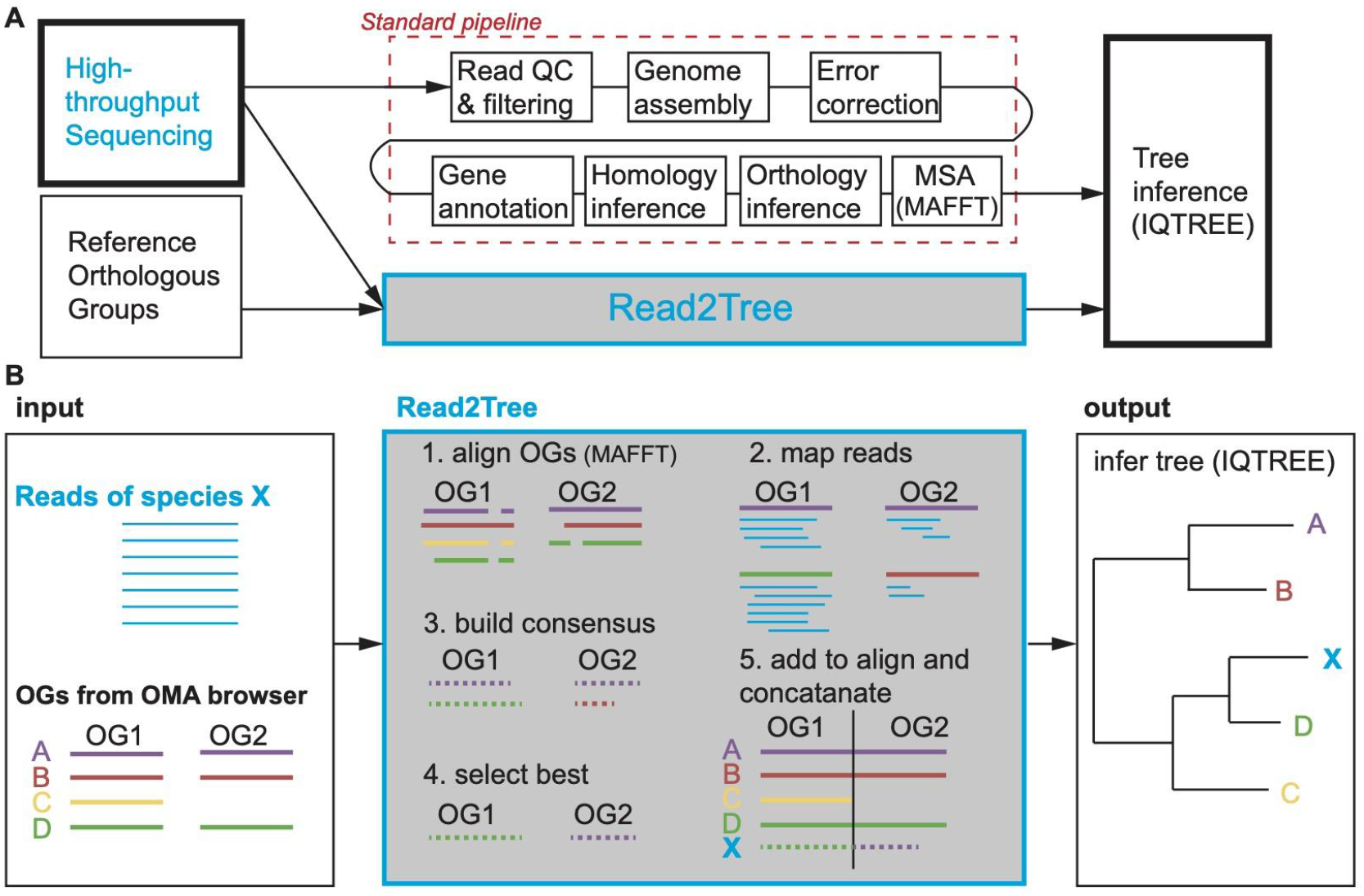
Strategy and pipeline explanation. (A) Read2Tree aims at sidestepping many time intensive and costly pipeline steps to obtain a phylogenetic tree when using many species, therefore going from read to tree. (B) Overview of the Read2Tree pipeline.

The current trend however is to sequence ever more species and samples. The Earth BioGenome Project, launched in Nov. 2018, aims at sequencing “all 1.5 million known animal, plant, protozoan and fungal species on Earth” within the coming decade^19^. The constituting consortia are making progress streamlining and optimising the sequencing and annotation process, but the orthology inference and tree inference steps remain highly challenging. In parallel, considerable genome sequencing activity is taking place in individual labs, with sample sizes of hundreds to thousands of genomes per study becoming common^16^. However, depending on the species of interest, high-quality reference genomes are often lacking, and individual labs often lack the computational infrastructure or expertise to fully leverage the data across individual analysis steps. This is exemplified in major consortia-led studies requiring years and millions of dollars to elucidate the evolution of certain species of interest. Or most recently, the use of various pipelines to assess variation and report assemblies from SARS-CoV-2. Thus a major bottleneck is becoming the harmonised analysis of these large-scale data sets to avoid certain biases or artefacts.

Here we introduce Read2Tree, a novel approach to infer species trees, which works by directly processing raw sequencing reads into groups of corresponding genes—bypassing genome assembly, annotation, or all-versus-all sequence comparisons. Read2Tree is able to provide a full phylogenetic comparison of hundreds of samples in a fraction of time compared to current established pipelines. Crucially, the speedup is achieved *without compromising the accuracy of the resulting trees*. In addition, Read2Tree is able to also provide accurate trees and species comparisons using only low coverage (0.1x) data sets as well as RNA vs. genomic sequencing and operates on long or short reads. This makes Read2Tree a highly versatile method to obtain key insight from a single sample scaling up to thousands of samples. To establish this novel approach we assess its performance on a battery of genomic and transcriptomic datasets spanning different kingdoms, divergence time, and sequencing technology. Subsequently we apply Read2Tree to construct a large yeast tree of life and apply it to compare SARS-CoV-2 samples—thus highlighting the accuracy (e.g. compared to NCBI classification) and speed of Read2Tree.

## RESULTS

State-of-the-art phylogenomic pipelines require many steps, which can be both time consuming and error-prone (**Fig 1A**). With Read2Tree, we directly process raw sequencing reads and reconstruct sequence alignments for conventional tree inference methods (**Fig 1B, SFig 1**). We start by aligning raw reads to nucleotide sequences derived from the genome-wide reference orthologous groups (OGs; we used Mafft^20^ as default) (**Fig 1B: 1**). Within each orthologous group, we reconstruct protein sequences from reads aligned to reference sequences (**Fig 1B: 2**). Importantly, these sequences in reference orthologous groups are not restricted to single-copy marker genes, such as the mitochondrial cytochrome *c* oxidase I (COI) gene or BUSCO genes ^21^; they also include multiple paralogous genes as well as non-universal genes. This is achieved by leveraging orthologous groups computed from 2500 diverse genomes analysed in the Orthologous Matrix (OMA) resource developed in our lab^22,23^. Next, we retain the best reference-guided reconstructed sequence, using the number of reconstructed nucleotide bases as criterion (**Fig. 1B: 3, SFig 2**). Subsequently, the selected consensus is added to the orthologous group’s multiple sequence alignment (**Fig 1B: 4**). Finally, putative orthologous group selection and tree inference can proceed using conventional methods (we use IQTREE^24^ by default; **Fig 1B: 5**). See Methods for greater detail on the individual steps.

This way Read2Tree is able to report key information across putative orthologous groups in a fraction of time over conventional comparative genomic pipelines—by bypassing genome assembly, annotation, homology, and orthology inference. Furthermore, because each sample is processed independently, Read2Tree can process the input genomes in parallel, and scales linearly with respect to the number of input genomes.

### Accuracy as a function of distance to closest reference and coverage

We tested Read2Tree on a wide array of conditions, with two kinds of sequence (DNA *vs*. RNA), three target species (*Arabidopsis thaliana, Saccharomyces cerevisiae* and *Mus musculus*), three types of sequencing technology (Illumina, PacBio and ONT), six levels of sequencing coverage (ranging from 0.2 to 20x), and six different sets of reference species (increasingly distant from the targets spanning over 1 billion years of evolution) (see **Fig 2A**). For sequence reconstruction accuracy (**Fig 2B**), we measured both the correctness of the reconstructed sequences (“precision”) and the completeness of the reconstructed sequences (“recall”). For tree reconstruction accuracy (**Fig 2C & SFig 6**), we compare the reconstructed tree with the known species phylogeny and report both the precision and the recall of the reconstructed trees, in terms of the branches with at least 90% support.

**Figure 2.**
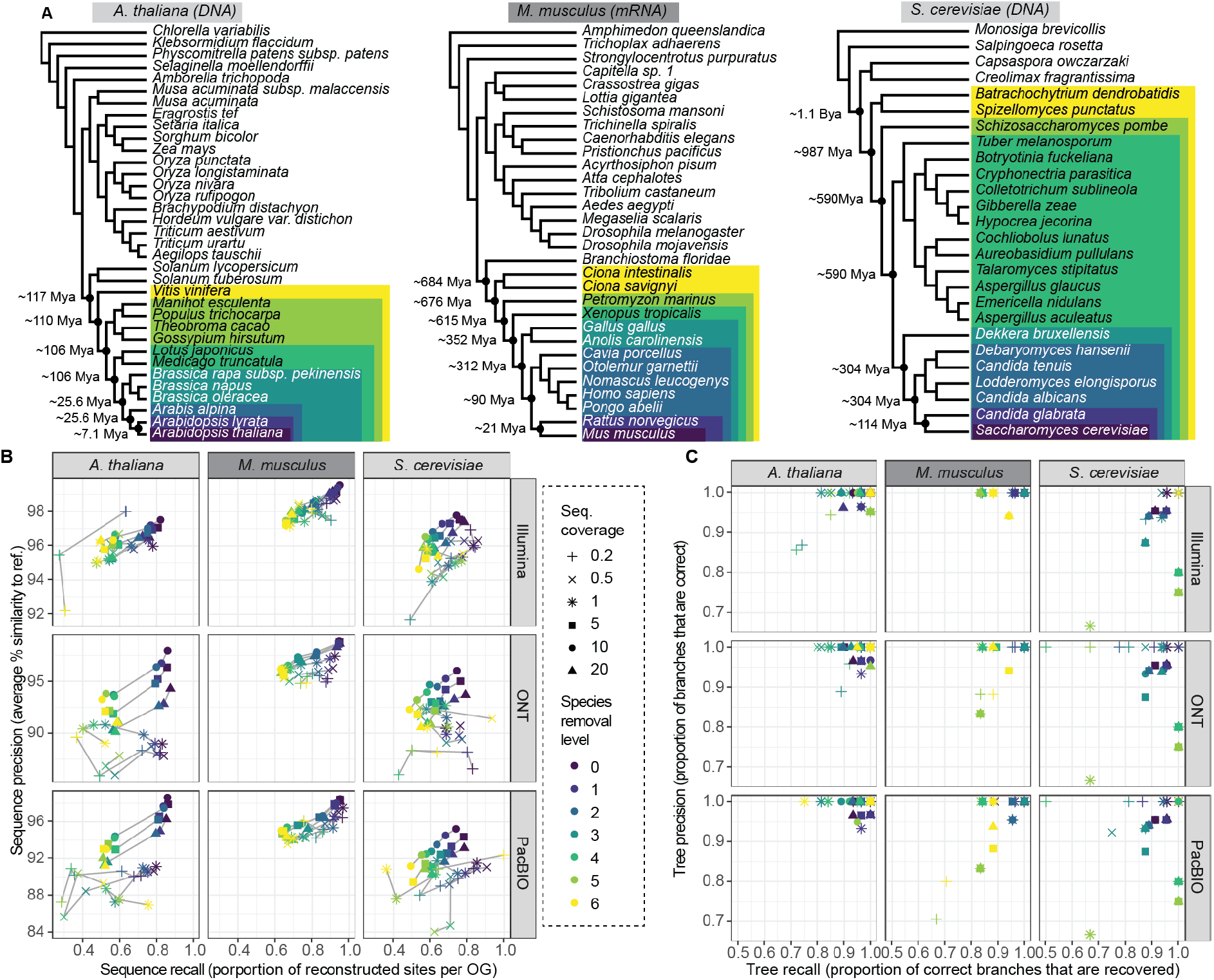
Benchmark of Read2Tree using three different datasets, six different coverage levels, and three sequencing technologies. (A) Phylogenetic trees of reference datasets. In dark purple (bottom) species used for mapping. Colours represent species removal to assess dependency on closest neighbours in reference datasets. Timepoints were obtained from timetree.org. (B) Read2Tree sequences are more similar (percentage identity) and more complete with increasing coverage and decreasing distance to a more closely related species. Best sequence identity is obtained for Illumina data. Colours convey the increasing evolutionary distance to the closest reference species. (C) Precision and recall of trees reconstructed using Read2Tree after collapsing branches below 90% support.

In general, Read2Tree was able to maintain a high precision in terms of sequence reconstruction (**Fig 2B**) and tree reconstruction (**Fig 2C**) across all datasets, with varying levels of recall depending on the dataset difficulty. First we assessed the effect of coverage ranging from 0.2x to 20x of the individual data sets. We observed that increasing the sequencing coverage had little impact on precision, and mainly lowered recall. Thus, remarkably, Read2Tree is able to maintain typically 90-95% precision at the sequence level even with coverages as low as 0.2x (**Fig 2B**). The best low-coverage results were obtained on transcriptomic short read data in mice, where precision reached 98.5% at 0.2x coverage. To assess the versatility of Read2Tree we benchmarked it across DNA and RNA data sets. This did not have a large impact in general, but transcriptomic RNA results (in the mouse dataset) are marginally less impacted by differences in average coverage, perhaps due to the large coverage variance from uneven gene expression levels in these data (see **Fig 2B&C**). Next, we assessed if Read2Tree is capable of utilising the range of current sequencing technologies. For this we applied it across traditional short reads, Oxford Nanopore and PacBio long reads. To enable this, Read2Tree has slightly different mapping strategies built in for long vs. short reads (see Methods). As **Fig 2B&C** shows Read2Tree maintained a high accuracy across each sequencing technology, but we observed the highest accuracy over traditional short reads. We have not assessed more recent sequencing technologies such as PacBio HiFi or Illumina infinity that might change this result.

Finally, we assessed the robustness of Read2Tree with respect to the evolutionary distance between the sample at hand and the closest relative in the reference set. This is often critical as one might not know the closest ancestor that is assembled or it is not available^25^. Thus we tested Read2Tree across a wide range of evolutionary distances ranging from 7 million years ago to over 1.1 billion years ago. While these are certainly extreme scenarios, overall Read2Tree was able to cope with them successfully. **Fig 2 B&C** show that the choice of reference set mainly impacted recall, with closer reference genomes leading to more reconstructed positions. Remarkably, Read2Tree was able to maintain high accuracy even in the datasets with very distant references—e.g. processing mouse RNA-seq data without any vertebrate genome in the reference set.

We also tested Read2Tree on simulated data, for coverages between 0.1x and 10x and distance to the closest reference varying between 2 and 150 point accepted mutation (PAM) units—where 100 PAM corresponds to 1 substitution per site on average. The reconstructed trees were perfect in all but the most extreme scenarios (PAM > 120 or coverage <0.5x; **SFig 7**).

Given the extensive benchmarks across species, coverage, sequencing technology, assay (DNA and RNA) and simulated data, we observe that Read2Tree is indeed a highly versatile and accurate tool to reconstruct phylogeny directly from raw reads.

### Read2Tree is faster and more accurate than assembly-based tree inference

Next, we compared the performances of Read2Tree with conventional assembly pipelines. For this we generated *de novo* assemblies and protein predictions across the same data sets as from the previous section, using Canu^26^ for PacBio and ONT data and Megahit^27^ together with SoapDeNovo^28^ for the Illumina reads (see methods). The conventional assemblies were processed using OMA standalone, including the same exported reference genomes, as OMA standalone was previously shown to identify the most accurate phylogenetic marker genes^29^. For the inclusion of orthologous markers in the concatenated alignment used for tree inference, we required a commonly set minimum threshold of 80% taxon presence. As above, we varied the closest remaining species in the dataset by removing species along the reference tree (**Fig 2. A**). With different coverages and reference sets, we obtained 42 data points per species. For each of these data points we performed the orthology inference separately and recorded its computation time. The proportion of sequences placed into the respective OGs showed high levels of variation (**SFig 8**). For each assembly and variation of proteomes, we computed the topological distance between the resulting tree from assembly or Read2Tree with trees obtained using high quality genome assemblies for *A. thaliana* and *S. cerevisiae*.

**Fig 3** shows the overall results highlighting the performance of Read2Tree. Perhaps unsurprisingly, we observed that coverage levels had a profound impact on the performance of assembly-based approaches rendering them incapable of dealing with coverages below 5-10x. Thus for these data sets we only report Read2Tree results.

**Figure 3.**
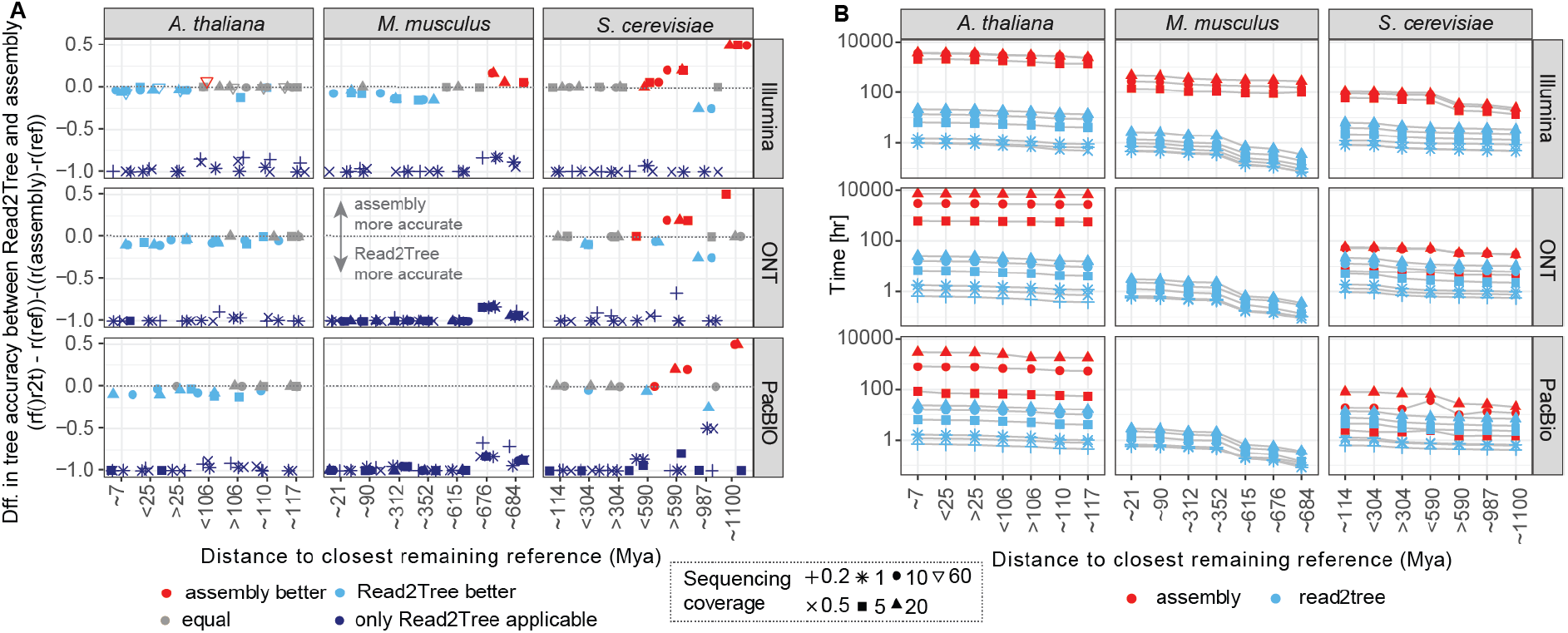
Comparison of Read2Tree with regular pipeline with assembly, orthology prediction and multiple sequence alignment computation. (A) Comparison of trees using the difference between the reference tree between either the tree of Read2Tree and the tree coming from the assembly approach. For dark blue we had only Read2Tree trees as assemblies for these low coverages are not possible to obtain. Below zero (in dark or light blue), Read2Tree is more accurate, while above zero (in red), the assembly approach is more accurate. In grey, no difference between the methodologies. (B) Comparison of wall time needed from reads to availability of concatenated multiple sequence alignment showing the dependencies of available closest remaining reference and coverage.

Where both approaches can be compared, the only cases where the conventional *de novo* assembly approach outperformed Read2Tree were with high coverage *and* very distant (>500 Mya) to the closest reference species (**Fig 3A**, upper right region of each graph). In all other scenarios, Read2Tree outperformed the conventional approach in accuracy. Specifically, on the yeast dataset at higher coverage level, both assembly and Read2Tree performed well overall—we never observed more than two different branches between the obtained and reference trees. With at least 10x coverage and distant reference species, the conventional assembly approach outperformed Read2Tree (**Fig 3A** and **SFig 4**).

By contrast, on the more complex *A. thaliana* and *M. musculus* datasets, Read2Tree outperformed the assembly approach—with fewer differences to the reference (up to 2 different branches for Read2Tree, versus up to 4 for the conventional approach). On the ONT data—characterised by longer reads but higher error rate, Read2Tree outperformed the conventional approach on both datasets.

Finally, in terms of compute time, Read2Tree was generally much faster than the conventional approach, up to 100x faster on the larger genomes (**Fig 3 B**).

Altogether, these results indicate that Read2Tree is faster in all conditions, and produces reliable trees in low coverage datasets and other datasets where the conventional approach fails entirely (long-read transcriptomics). At higher coverage levels, the trees inferred by Read2Tree rival in quality with those obtained from assembled reference species with a full pipeline, particularly when applied to more complex genomes, and unless the closest reference species is very distant (> 500 million years).

We also compared Read2Tree with MASH, a fast *k*-mer-based approach^30^ commonly used on bacterial genomes. While the alignment-free approach of MASH was much faster than even Read2Tree, the resulting trees were much less accurate than either Read2Tree or the assembly-based approach (**SFig 5**). This illustrates why alignment-free approaches such as MASH, while very useful for fast approximations, are typically not suitable to reconstruct high-quality phylogenetic trees.

### Read2Tree accurately reconstructs a yeast tree of life encompassing 435 genomes

To assess a potential large scale application for Read2Tree, we applied it to reconstruct a large yeast phylogeny from raw reads. Thanks to Read2Tree’s ability to process low coverage datasets, we could extend our analysis to all Illumina single and paired-end, ONT, PacBio and 454 Sequencing read datasets available for budding yeast in the NCBI SRA database (November 2018, 404 species) and 31 reference species obtained from the OMA Database (release 2018, 3063 OGs). Using an automated approach for retrieval and mapping, we were able to obtain direct sequences for 404 species (2019, Supplementary File 1). Read2Tree could process these datasets in around a month of computation (adding each species sequentially and performing the mapping on 30 CPUs — one CPU per reference — in parallel), due to its “embarrassingly parallel” architecture–with every sample being processed independently up to phylogenetic inference (10x Illumina: ~20 minutes using 4 threads).

A large proportion of these data sets were recently used to construct a phylogeny across 363 budding yeast species^31^. This included a dataset of 196 new assemblies and their annotations^31^. This large effort provided the first delineation of the yeast tree of life into 13 main clades and highlighted the influence of horizontal gene transfer in the evolution of yeast species^31^. Due to the complexity of state-of-the-art pipelines, it also consumed millions of CPU hours and years of work. Furthermore, the conventional assembly-based approach could not include low-coverage samples into their analysis. We were able to extend this work using Read2Tree using a fraction of the resources.

Using Read2Tree we were able to compute and produce this large phylogeny across 435 samples (including 31 species as reference). Some of the samples failed due to their too low coverage levels of around 3.1x assuming a 12Mbp long average genome size. Nevertheless, using Read2Tree we were able to include multiple samples even at coverage levels below 5x which were reported with over 2500 sequences placed in orthologous groups (**SFig 14**). Read2Tree was able to reconstruct the phylogeny and also reported the phylogeny relevant genes assembled per sample which overall showed similar GC levels as the reference data (**SFig 15**). This was also exemplified by the fact that we did not observe a correlation between the number of sequences placed into OGs per species and their individual coverage (**SFig 14**, correlation 0.2).

Considering the subset of species in common, our results were highly congruent with those of Shen *et al* ^31^ (**Fig 4, SFig 12 and SFig 13**): both trees exhibited similar distances to the NCBI taxonomy tree—297 ours vs. 291 Shen *et al* splits respectively. In direct comparison, Shen et al. and Read2Tree were more similar with one another, with only 128 different splits (20% difference of the branches), than either was to the NCBI taxonomy. After collapsing branches with a support below 90, the difference in the number of splits between the conservative NCBI tree and ours was 29 splits and between Shen *et al* 25 splits. Twenty-four of these splits were in common between Read2Tree and Shen *et al*. To get more insights on the nature of these differences, we assessed the agreement with the NCBI taxonomy for two different levels of resolution: family and genus. At the coarser family level, Read2Tree was more consistent with the NCBI taxonomy for six families, while Shen *et al*. was more consistent in one family (**SFig 10**). At the finer genus level, Read2Tree was more consistent with the NCBI taxonomy for four genera, versus ten for Shen *et al*. (**SFig 11**).

**Figure 4.**
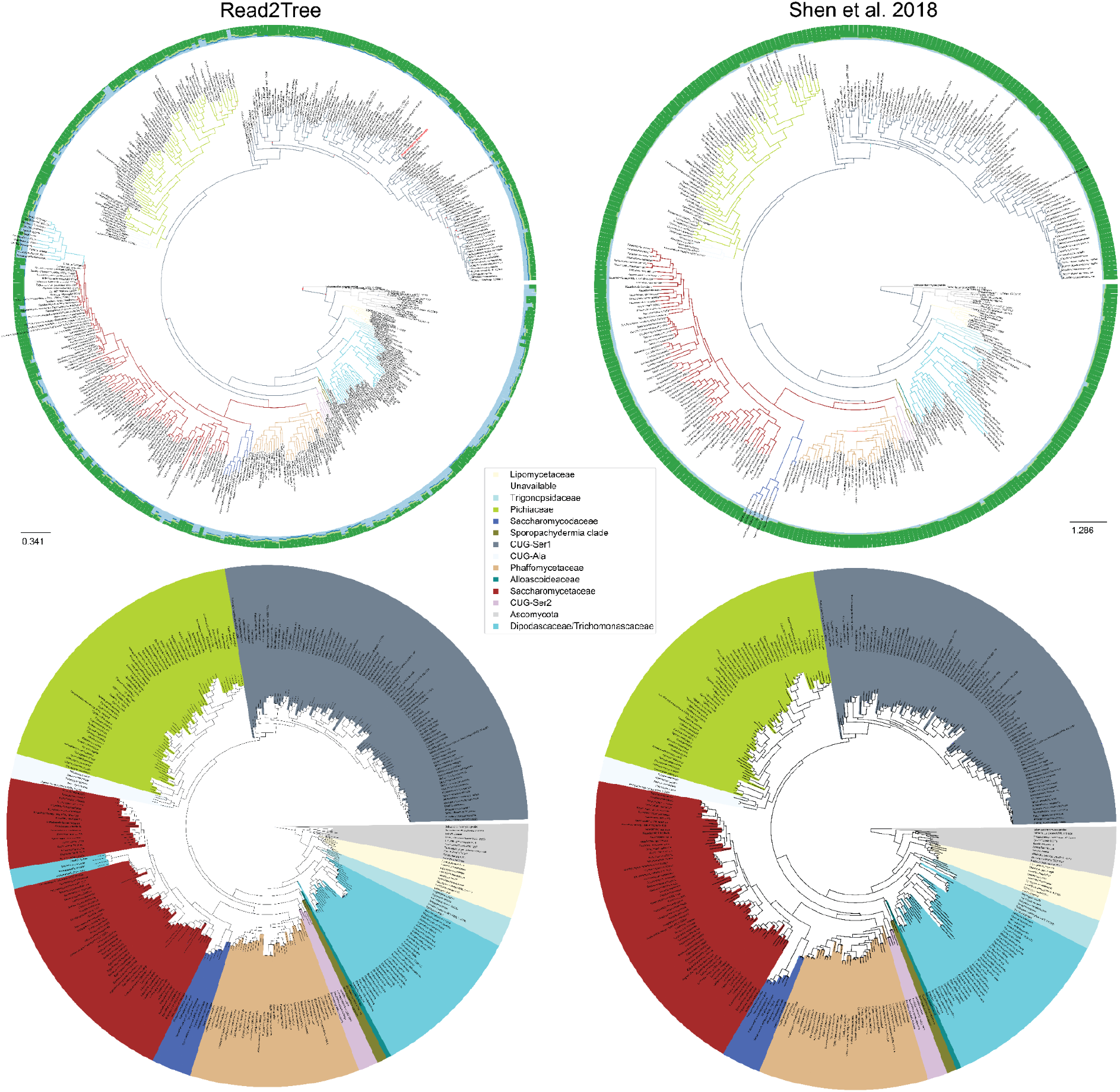
High consistency between Read2Tree and a state-of-the-art phylogenetic pipeline^31^. The top row shows full trees and the alignment matrix used to compute the tree as outer circles. Red dots indicate nodes with bootstrap below 100. The species Naumovozyma dairenensis, previously misclassified^32,33^, is highlighted in red. Bottom row shows trees trimmed to an overlapping leaf set.

Nevertheless, there are still certain differences between Read2Tree and the NCBI taxonomy remaining. While resolving most such instances would constitute entire follow-up studies in their own right, we were able to explain one apparent disagreement: *Naumovozyma dairenensis* is placed in the CUG-Ser1 classification, while according to the NCBI taxonomy, it should be an ascomycetous yeast in the Saccharomyces sensu lato group within the family Saccharomycetaceae. However, this is a case of erroneous metadata reported in the literature.^32,33^

Given this phylogeny, we can now easily update and extend it using Read2Tree in a matter of minutes with additional sequences being generated. This enables a deep dive in the comparative genomics of yeast and explore further their differences between the strains and their impact on live, food production etc. This is also easily reproducible for other organisms as Read2Tree is capable to span large evolutionary distances with respect to the reference tree.

### Read2Tree for zoonotic surveillance and human epidemiology

To further illustrate the versatility of Read2Tree, we used it to reconstruct a phylogeny encompassing various coronaviruses from the OMA coronavirus database as well as 215 raw coronavirus sequencing samples deposited to the Short Read Archive. Besides the putative SARS-CoV-2 sequence, we also included two samples from bat (SRR11085797^34^ and SRR11085736^35^), and one from mink^36^ (SRX9605666).

The reconstructed phylogeny was in complete agreement with the lineage classification obtained from the UniProt reference proteomes. In particular, the tree not only recovered the main coronavirus genera (*Alpha*-, *Beta*-, *Gamma*-, and *Deltacoronavirus*), but also all subgenera with complete consistency (**Fig 5**).

**Figure 5.**
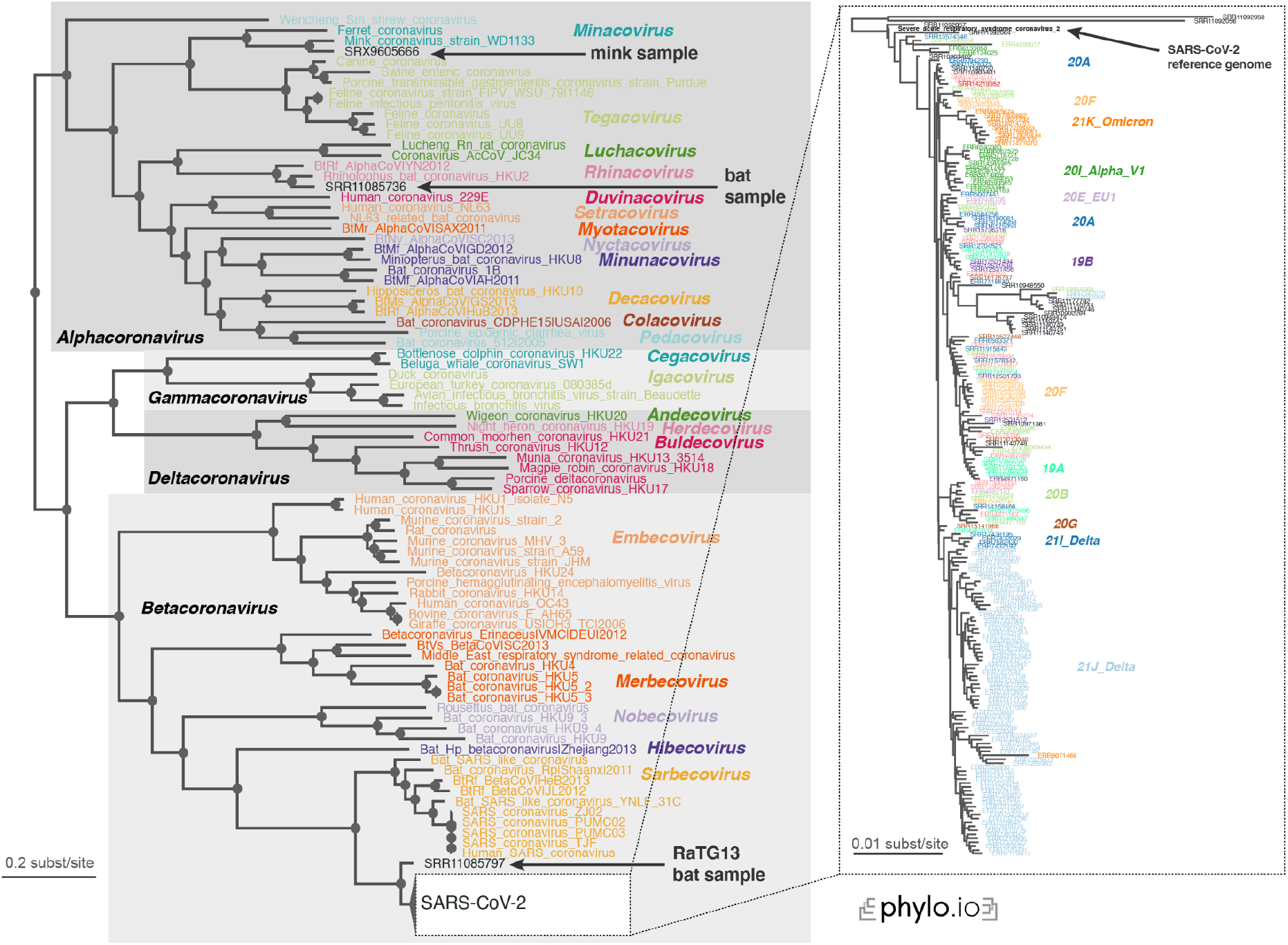
Read2Tree correctly classifies the recent SARS-CoV-2 sequences and recapitulates the evolution of the individual variants. All genera (grey boxes in the overall tree) and subgenera (coloured labels) are correctly delineated. The inset focuses on the part of the tree with 215 SARS-CoV-2 samples, and variants of concern (coloured labels) cluster consistently on the tree, indicating that Read2Tree can be used to categorise the samples.

The first bat sample corresponds to the reads of RaTG13, which is the closest relative of *SARS-CoV-2* identified yet^34^. Indeed, in our tree it falls right outside the *SARS-CoV-2* clade. The other bat sample as well could be confirmed as an *Alphacoronavirus*, subgenus *Rhinacovirus^35^*. Likewise, we could confirm the classification of the mink sample, identified as an *Alphacoronavirus*, subgenus *Minacovirus* by the authors^36^.

The position of the *SARS-CoV-2* sequences within the coronavirus tree of life is also consistent with our prior knowledge on them. The reference genome, the Wuhan-Hu-1 sequence reported in early January 2020^37^, is at the base of the subtree. The only three sequences that branch out prior to it are SRR11092056-8—which were obtained from patients with severe pneumonia at the beginning of the pandemic^34^. Finally, we note that the variants of concern included in the analyses appear clearly as distinct clades on the tree.

To empirically test the scalability of our method, we also used Read2Tree to process 10,283 SARS-CoV-2 samples. The reconstructed tree clustered the sequences according to CDC variants of concerns classification, providing further evidence that the tool can be used to quickly and reliably classify SARS-CoV-2 variants (**SFig 17**). The same observation held for additional controls—running Read2Tree using coding-genes markers only (**SFig 16**), and using FastTree^38^ as tree inference method (**SFig 18**).

Overall, this application of Read2Tree to diverse coronaviruses sequences illustrates the ability of the tool to deal both with the considerable phylogenetic breath of this family of virus^39^ and the depth required to classify individual SARS-CoV-2 variants of concerns. This makes Read2Tree suitable for both zoonotic surveillance as well as human epidemiology^40^.

## Discussion

We presented Read2Tree, a novel approach to scale and ease the laborious process of comparative genomics: assembly, annotation, phylogenetic comparison. These steps are computationally costly, error-prone and require specialised knowledge. Using Read2Tree, we can directly reconstruct phylogenetic relevant genes from raw reads and thus enable a placement and comparison of the species at hand with minimum compute and coverage requirements. The efficiency of the approach makes it possible to process a large number of samples in parallel, using a consistent methodology, and without compromising accuracy compared with state-of-the-art pipelines.

Current inherent problems of large scale comparative genomics or in general comparative genomics projects recently shifted from obtaining accurate assemblies to annotation and curation of these assemblies. This was in part possible due to sequencing technology advancements over long reads^16,18^, but also due to innovations in assembly algorithms^41,42^ . These steps still require high DNA quality and are in general more expensive, but enable large projects such as the Vertebrate Genome Project^43^, the human pangenome^44^ and telomere-to-telomere^45^ projects. Nevertheless, in every of these cases the annotation of the genomes and the improvements in terms of continuity and accuracy remain major bottlenecks. Using Read2Tree these limitations can be overcome even with low-coverage, cost-effective Illumina data. Indeed, we showed that Read2Tree enables accurate analysis across all three sequencing technologies (Illumina, ONT and PacBio). All this can be achieved in a fraction of time and computational resources, thereby contributing to bringing large-scale phylogenomics within the reach of individual laboratories. Furthermore, large scale consortia could also benefit from running Read2Tree, despite having high coverage data sets, to independently QC their assembly and tree building approaches. Here Read2Tree can provide valuable insights within minutes.

One major advantage is that despite side-stepping de novo assembly, Read2Tree can operate in the absence of close reference genomes; indeed we demonstrated accurate tree reconstruction involving sequencing reads from species separated by hundreds of millions of years of divergence. Though we also reached some limits to this robustness, when subjecting Read2Tree to both very high divergence and low sequencing coverage, it should be noted that evolutionary distances will tend to diminish as ever more species get sequenced across the tree of life.

Furthermore, while most authors of genome resources deposit annotation sets alongside the assembled sequences, not all of them do. The ability to process genomes directly from raw reads not only circumvents this limitation; it can reduce the biases arising from overreliance on specific reference genomes, typically model organisms for which genomic resources tend to be more developed. There have been some initial efforts to “dehumanise” non-human great ape genomes^46^, but many other clades still suffer from analogous biases, which can be greatly reduced by processing raw reads.

We demonstrated the speed and accuracy of Read2Tree over a large scale yeast data set. Here, Read2Tree was able to reconstruct a high-quality tree from raw read samples directly retrieved from the Short Read Archive. This was achieved despite orders of magnitudes of variation in the coverage levels and other properties of the data sets (e.g. reads were generated across 10 years of sequencing, on a diverse set of sequencing instruments).

In a second illustrative application, we reconstructed a tree from raw coronavirus sequencing data, including 10 thousands samples from the ongoing SARS-CoV-2 pandemic. Here Read2Tree was again able to classify and place all samples correctly, be it across the full breadth of the *Coronaviridae* genus, or across the depth of minute variations among SARS-CoV-2 samples. This level of performance is remarkable, because the optimal choice of phylogenetic marker genes typically depends on the level of sequence divergence^47^.

We also compared Read2Tree with an ultrafast, alignment-free approach (MASH) where Read2Tree achieved a much higher accuracy (**SFig 5**). In its current form, Read2Tree serves a distinct function from metagenomic classifiers such as Kraken2^48^ or Centrifuge^49^. Indeed, while these tools seek to exploit known characteristic sequences for read-level taxonomic classification, Read2Tree aims at efficiently extracting the genome-wide (or transcriptome-wide) phylogenetic signal by inferring large multilocus input data matrices for phylogenetic tree inference tools, a step which has been shown to be critical to resolve difficult phylogenies^17,29,50–52^. Nevertheless, Read2Tree could be further developed to process metagenomic samples—by combining it with a genome binning pre-processing step. In recent years, a number of different approaches for genome binning have been proposed, be it through “differential coverage” approaches, which exploit correlated abundance across samples to identify reads coming from the same species^53–55^, using Hi-C protocols, which make it possible to identify parts of DNA in close physical proximity^56,57^, or single-cell technologies^58^.

Overall Read2Tree is a novel approach reconstructing phylogenetic important genes and characterising the sample at hand or entire sample collections, thereby enabling the study of a large number of genes and their evolution with no preprocessing, few computational resources, and minimal bioinformatic expertise. This will hopefully enable faster and more comprehensive phylogenetic reconstruction efforts—from tiny virus genomes to large eukaryotic ones, but also cell lineage, cancer trees, and other kinds of phylogenies across biology and medicine.

## METHODS

### Description of Read2Tree method

Read2Tree incorporates various publicly available tools for some of its steps (MAFFT^20^, NextGenMap^59^, Samtools^60^) and uses these in a structured manner to go from reads and reference OGs to a concatenated alignment that is fed directly into a tree inference tool, which by default is IQTREE^24^. For this purpose, it needs two sets of input data: i) a set of reference orthologous groups that can be obtained directly from the OMA database; ii) the reads to be mapped coming from a single species. The Read2Tree pipeline works in the following way. First, it retrieves DNA sequences using the REST-API from the OMA-browser from the selected reference orthologous groups, then sorts these into one file per species. In parallel, it computes alignments using the AA sequence with MAFFT^20^ and then uses the Codon information to generate DNA alignments. Once computed, all reads are mapped against the DNA reference species and a consensus sequence is constructed (local assembly). Since our local assemblies are reference guided they can never be longer or shorter than the longest or shortest sequence part of an OG. Local assemblies are then placed into the alignments using the coordinates of the best selected reference. Therefore, no new alignment is necessary and we can assure that the right AA / DNA is placed in the right position in the alignment. The resulting alignments for each OG are then concatenated and a tree is computed. More details about the inner workings of Read2Tree are provided in **SFig 1**.

Read2tree can be parallelized using multiple instances across the mapping step. It is recommended to compute the reference set first. The mapping step can then be split such that each mapping can be performed as single job submission on high-performance clusters. Read2Tree was developed in python. Read2Tree is open source (MIT License) and available online at https://github.com/DessimozLab/read2tree.

### OG selection

Orthologous groups were selected from OMA^61^ using the marker gene export functionality (https://omabrowser.org/oma/export_markers/). For all species the maximum number of covered species was set to 0.8 and maximum number of markers to −1 (unlimited). Species selected are displayed in **Fig 1A**.

### Reads

Whole genome sequencing reads for *A. thaliana* and *S. cerevisiae* were obtained from the SRA database for technologies PacBio, Illumina and Oxford Nanopore. mRNA sequencing reads for *M. musculus* were also obtained for all three technologies from the SRA database. Subsampling of reads was performed in python (see repository). For PacBio and ONT reads subsampling was optimised such that cumulative number of bases fits to the expected coverage. For coverage test reads were subsampled assuming for mouse 38 Mbp accumulated gene length (transcriptome), thale cress 120 Mbp and yeast 12 Mbp genome lengths. Reads were sampled to obtain 20X, 10X, 5X, 1X, 0.5X and 0.2X coverage levels. Reads for the big yeast tree were obtained from the SRA database (Supplementary File 1). Reads for coronavirus were obtained from the SRA database (Supplementary File 1). All SRA numbers are available in Supplementary File 1.

### Reference tree construction

Reference trees for the 3 evaluated species were computed using the species as defined in **Fig 2A**. Species were selected from OMA ^61^ as described in the OG selection. Individual OGs (gene markers) were aligned using MAFFT^20^ v7.310 (--maxiter 1000 --local) and trees were inferred with IQTREE^24^ v1.6.9 (-m LG -nt 4 -mem 4G -seed 12345 -bb 1000). For reference trees that were used for testing the dependency on the reference dataset, specific species were deleted from existing alignments and trees were computed with IQTREE as stated before. All reference trees are available in the Supplementary File 2. To highlight the years of evolution, we collected the time using timetree^62^ (April 2022).

### Read2Tree runs

For the single species runs part of **Fig 2 & 3**, Read2Tree was run with default parameters. For the large yeast tree **Fig 4** Read2Tree was run in multiple steps. First the reference dataset was obtained (using the --reference option). Then mapping was parallelized such that for each species the mapping against a single reference was performed individually (using --single_mapping option). This means that for each species 31 parallelized mappings were performed. Additionally, species with reads with bigger than 20X coverage were sampled to 20X coverage assuming a genome length of 12Mbp. Subsampling of reads is integrated into the Read2Tree workflow. Finally, all mapped species were merged together and concatenated to provide the multiple sequence output (using --merge_all_mappings).

### Accuracy assessment

We assessed the accuracy of sequence reconstruction by taking each Read2Tree reconstructed sequence (for each species, coverage, technology and removal level) placed in an OG and performed a blastp (ncbi-blast; v2.8.1) search against its original OG that contained the original sequence coming from a high quality assembly for the species of interest. Accuracy was measured as blast percentage identity and recall as the total number of obtained amino acids in the concatenated Multiple Sequence Alignment (MSA) of all OGs. Additionally, we evaluated whether the top hit of the Read2Tree reconstructed sequence was most similar to its assembled same species counterpart, or the sequence used as reference for reconstruction or any other random sequence part of that particular orthologous group (**SFig 3**).

### Assemblies

For the three species, the whole genome data was assembled with individual sequencing technology specific assembly programs following best practice or default parameters. For Illumina we first used megahit ^27^(v1.2.9) with default parameters for assembling the contigs. Subsequently SOAPdenovo^28^ (version 2.04-r241) for scaffolding: First, SOAPdenovo-fusion -D -K 41 -c megahit.contigs.fa -g scaffold_prefix -p 20 followed by SOAPdenovo-63mer map and scaff with recommended parameters over the config file. For ONT reads we assembled the reads using Canu^26^ (v2.0) with a specified genome size (genomeSize) gnuplotTested=true -nanopore-raw and useGrid=false parameters to run it locally on only one node on the cluster. Lastly for PacBio CLR data we also used Canu (v2.0) with similar parameters, but specifying the -pacbio-raw parameter. All runtimes were measured using linux time and the wall and CPU Time were recorded. The RNA seq data was assembled differently to the whole genome. For Illumina RNA seq, we used Trinity^63^ (v2.8.5) with the following parameters: --seqType fq --max_memory 50G --left reads1.fq.gz --right reads2.fq.gz --CPU 6 --trimmomatic --full_cleanup --output prefix. These execute Trimmomatic automatically and follow the recommendations from trinity.

### Orthology prediction of assembled genomes

For each assembly (species, technology and coverage level) we run OMA standalone (v2.3.3) on the UNIL HPC clusters using a SLURM scheduler. For this we collected all the species as depicted in **Fig 2** using the OMA All vs. All export function. Then we removed according to **Fig 2** the relevant species, adding each time the assembly for mouse, yeast or thale cress in the set and run the orthology prediction with standard parameters (OMA v.2.2.1). Thus for instance for the Illumina *M. musculus* 10X assembly we run OMA 7 times for all reference datasets with increasing distance to its closest relative. In total we run 126 different OMA runs with 7 variations of reference proteomes and 3 variations of technologies 3 coverage levels for *A. thaliana* and *S. cervisiea*. Additionally, we run 21 times OMA for M. musculus for 5X, 10X and 20X Illumina assemblies. The all vs. all part was parallelized on 1000 nodes and the final part was run on a single node with 40G memory. To obtain OGs for tree inference we applied the 0.8 taxonomic occupancy threshold as previously. OGs were filtered according to the procedure in Shen et al. (see below). OGs were individually aligned using MAFFT^20^ v7.310 (--maxiter 1000 --local), concatenated and trees were inferred with IQTREE^24^ v1.6.9 (-m LG -nt 4 -mem 4G -seed 12345 -bb 1000).

### Tree vs. tree comparison

Each Read2Tree tree was compared to a fitting reference using several tree distance measures. For topological similarity we used two approaches, one that uses the Robinson-Foulds distance and counts the number of different splits between two trees and one that collapses each node with a bootstrap support below a certain threshold and then counts the number of overlapping splits. Then we define as recall the number of overlapping splits divided by the number of splits in the reference and as precision the number of overlapping splits divided by the number of splits in the Read2Tree tree.

### Large yeast tree

For the large yeast tree we extracted all yeast available datasets in the SRA November 2018 (406 species, Supplementary File 1) and applied Read2Tree (standard parameters) to 31 yeast species extracted from the OMA database (Nov 2018) using the marker export function with min tax availability of 0.8 (3082 OGs). Selected species are available in the Supplementary File 3. Reads from the SRA database were mapped according to their sequencing methodology using Read2Tree. In order to compare our analysis to Shen et al 2018 we aimed to have as many species in common as possible. For this purpose, we complemented our tree with additional sequences that we simulated from missing species in our tree that were present in the Shen et al 2018 tree. Simulations were conducted with iss (v1.3.0 https://github.com/HadrienG/InSilicoSeq, --model hiseq -n 600000). In order to map the species from the Shen *et al*^31^ tree to ours, we obtained the taxon id of species / strains using ete3’s NCBI interface^64^. For species where automated mapping was not possible we obtained the taxid using the NCBI taxonomy interface (https://www.ncbi.nlm.nih.gov/Taxonomy/Browser/wwwtax.cgi).

### Filtering OGs yeast as in Shen *et al* 2018

Given the reconstructed sequences placed in their respective OGs and added to their alignment, we decided to compute a tree following the protocol of ^31^. In brief, from the 3082 alignments we selected the ones that contained more than 171 species resulting in 1829 OGs. Then we used phyutils 2.2.6 (seqs -aa -clean 0.01) to clean up the alignments. Since our approach does not place multiple sequences from the same species into one OG, we skipped the removal of putative paralogs. Within the alignments we changed all “X” with the gapcharacter “-”. Then we applied trimAl v1.4.rev15 (-gappyout). Then we removed protein sequences whose lengths were shorter than 50% the length of the trimmed multiple sequence alignment length of each OG they belonged to. We also removed OGs whose total trimmed multiple sequence alignment length was < 167 amino acid sites. These resulted in 926 alignments. With these alignments we used IQTREE (v1.6.9) with automatic model selection to compute trees. Then we identified species in the gene trees that had a branch length longer than 20 times the median of all branch lengths. We removed these species from the respected alignments again controlling that the more than 171 species are included. Then we computed the tree using IQTREE (-seed 12345, -m LG+G4, -bb 1000, -nt 20).

### Large yeast tree comparison

Using all taxon ids, we retrieved the current uptodate NCBI reference taxonomy and the classification of each species. We then compared the three trees (NCBI, Read2Tree and Shen *et al*^31^) using the Robinson-Foulds distance on the overlapping leafset. Additionally, we overlaid the Shen *et al*. classification on our tree. Finally, we compared the trees using the ancestral node that contains the highest number of monophyletic species given a specific grouping (order, family, phylum) extracted from the NCBI taxonomy information. All comparisons were conducted using custom python jupyter notebooks. Additionally, we collected data on GC content, input coverage to mapping ratio. Trees were visualised with ete3^64^. The tanglegram plot was produced using the dendextend R library ^65^. Side by side topological comparison was obtained using phylo.io ^66^.

### *Coronaviridae* tree reconstruction

Marker genes were exported from https://corona.omabrowser.org/ with at least four species. DNA sequences for these genes were obtained from the same resource. Four extra groups with intergenic regions from the SARS-CoV-2 reference genome were added using a custom script. We extracted consecutive chunks of at least 30 bp from the reference genome MN908947 assembly that were not covered by either any CDS region or proteins not belonging to any OMA group in the https://corona.omabrowser.org resource (i.e. ORF8 and ORF10). This led to 4 regions (1..265; 26473..26522; 27760..27893; 29675..29903) which we treated as additional groups. SARS-CoV-2 samples were obtained from Nextstrain open (https://data.nextstrain.org/files/ncov/open/global/metadata.tsv.xz)^7^. Different samples with SRA accession that span all different clades were obtained with a custom python script (included in the linked repository below). SRA read accessions together with the clade annotations from nextstrain are available in Supplementary File 1. Reads were downloaded from the SRA database and trimmed. Read2Tree was applied to this dataset and all obtained reads were mapped to the marker genes. Read2Tree was run with standard parameters. The resulting supermatrix alignment was filtered by removing columns that had more than 70% gaps. This removed 30969 columns resulting in a supermatrix of size 295 x 42669. Finally, the tree was inferred using IQTree2 ^24^ (version 2.2.0-beta) with parameters -m GTR -ninit 2 -me 0.05.

As additional controls, we computed the trees with FastTree^38^ version 2.1.11 instead of IQTREE2 and without the additional four extra groups. All trees are available in Supplementary File 2.

For the scale up experiment with 10,283 samples, we used the same protocol, except for the source of the read annotations. Here, we used the clade annotations from https://harvestvariants.info/ (Accessions and annotations available in Supplementary File 1).

### Simulated phylogeny analysis

The simulated phylogeny includes a fixed topology for species tree with 15 species using the ALF package^67^ (version v0.99). We varied the branch length leading to one of the species (species of interest) between 2 PAM and 150 PAM. For each run, we infer afterwards the OMA Groups (excluding the species of interest). Then using art_illumina^68^, we generated DNA sequencing reads (paired-end) with length of 100 and 150 and coverage of 0.1 to 10. Next, for each case, we ran Read2Tree to infer the phylogeny. Finally, we calculated the Robinson–Foulds metric between inferred species tree and the true one based on the output of ALF.

### Comparison with MASH

We took established assemblies as a reference that we downloaded from NCBI. Subsequently, we used MASH (version 2.3) sketch^30^ with a size of 10m (k=21 as default) which was followed by MASH distance to obtain distances between the genomes and analysed the reads against that reference set. Finally, we applied RapidNJ ^69^ (version 2.3.2) on the distance matrix obtained from MASH to infer the species tree. We did that for different distances across the references that were provided, always comparing the reads from e.g. *A. thaliana* to the assemblies.

## Supporting information

Previous supplement

## Contributions

CD and FJS designed the study. DD and FJS implemented the software. DD, FJS, AA, SM performed data analysis and code review. DD, FJS, and CD drafted the manuscript. All authors edited and approved the manuscript.

## Acknowledgments

FJS is supported by NIH grants (UM1HG008898) and the National Institute of Allergy and Infectious Diseases (1U19AI144297). DD, SM, and CD were supported by Swiss National Science Foundations grants 183723, 186397 and 205085 (to CD).

## Data availability

References used and all SRA numbers of reads used are available in the supplement.

Supplement, scripts and reference data are available at https://github.com/dvdylus/read2tree_paper.git.

## Code availability

The source code for Read2Tree is available at https://github.com/DessimozLab/read2tree under an MIT open source licence.

## Read2Tree — Supplementary Figures

**Supplementary Figure 1.**
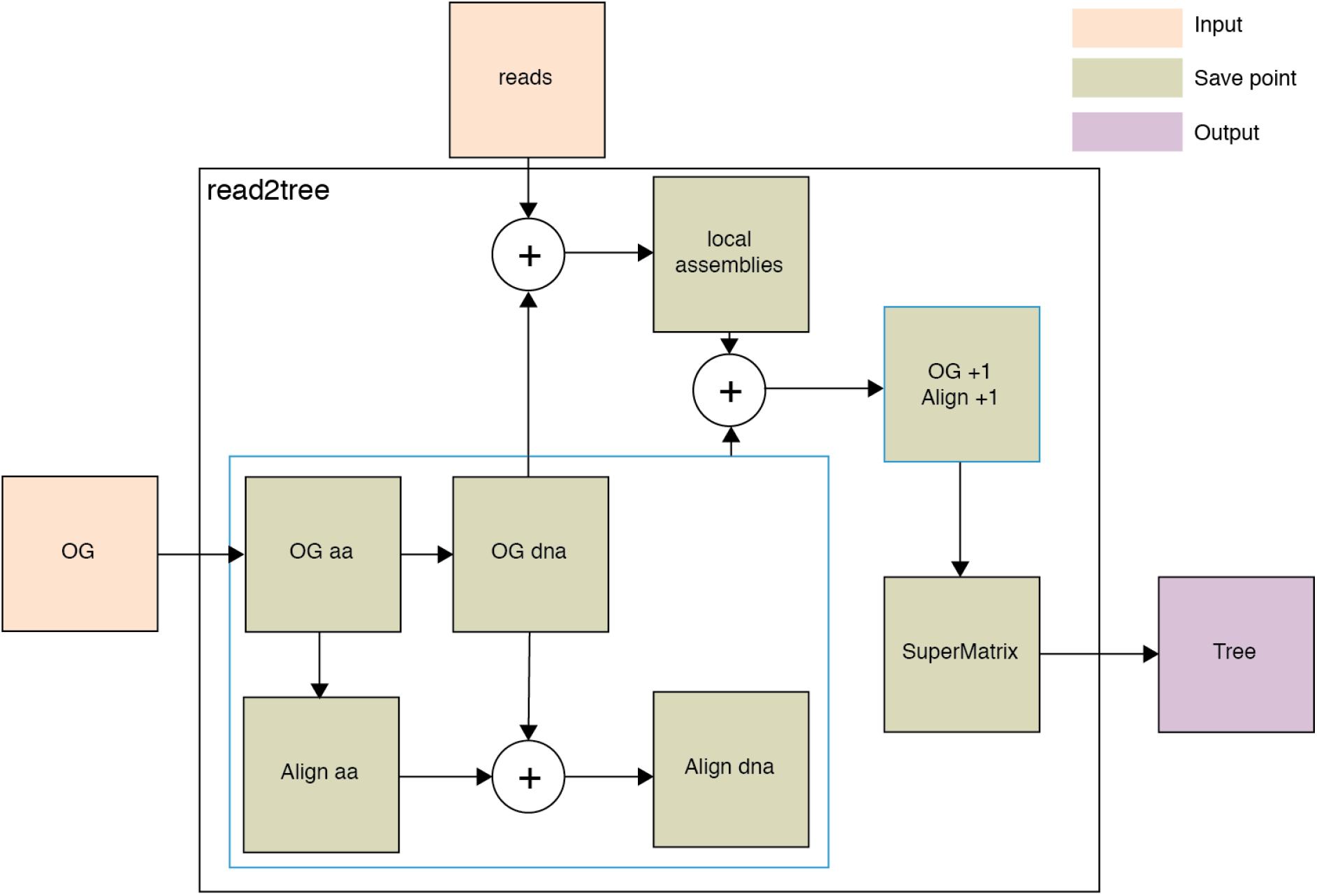
Graphical representation of pipeline. All boxes in green are stored by Read2Tree. Inputs are reads and a set of reference orthologous groups that can be selected from over 2000 species from the OMA database. Local assemblies here as reconstructed sequences using the bases placed against the reference.

**Supplementary Figure 2.**
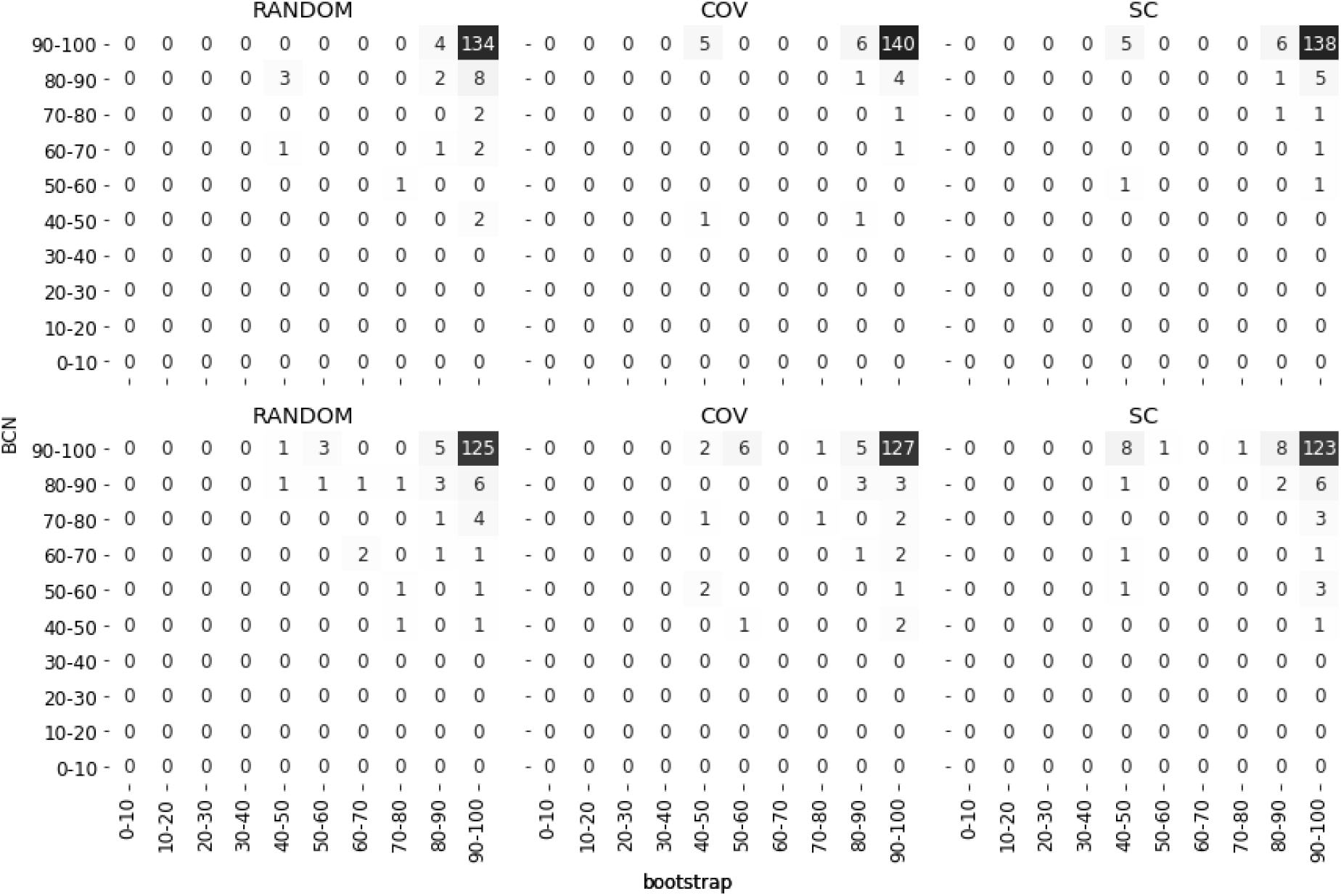
Comparison of different gene selection methods in 20 times 8out/8in test show differences in relation between bootstrap on nodes of mapped species and its jaccard similarity with reference tree (Best Corresponding Node BCN) value. Top row shows values obtained using simulated reads and bottom row shows values obtained using real reads. In both cases we see that using coverage as a selection method shows the highest number of nodes where mapped species have high bootstrap and high BCN values.

**Supplementary Figure 3.**
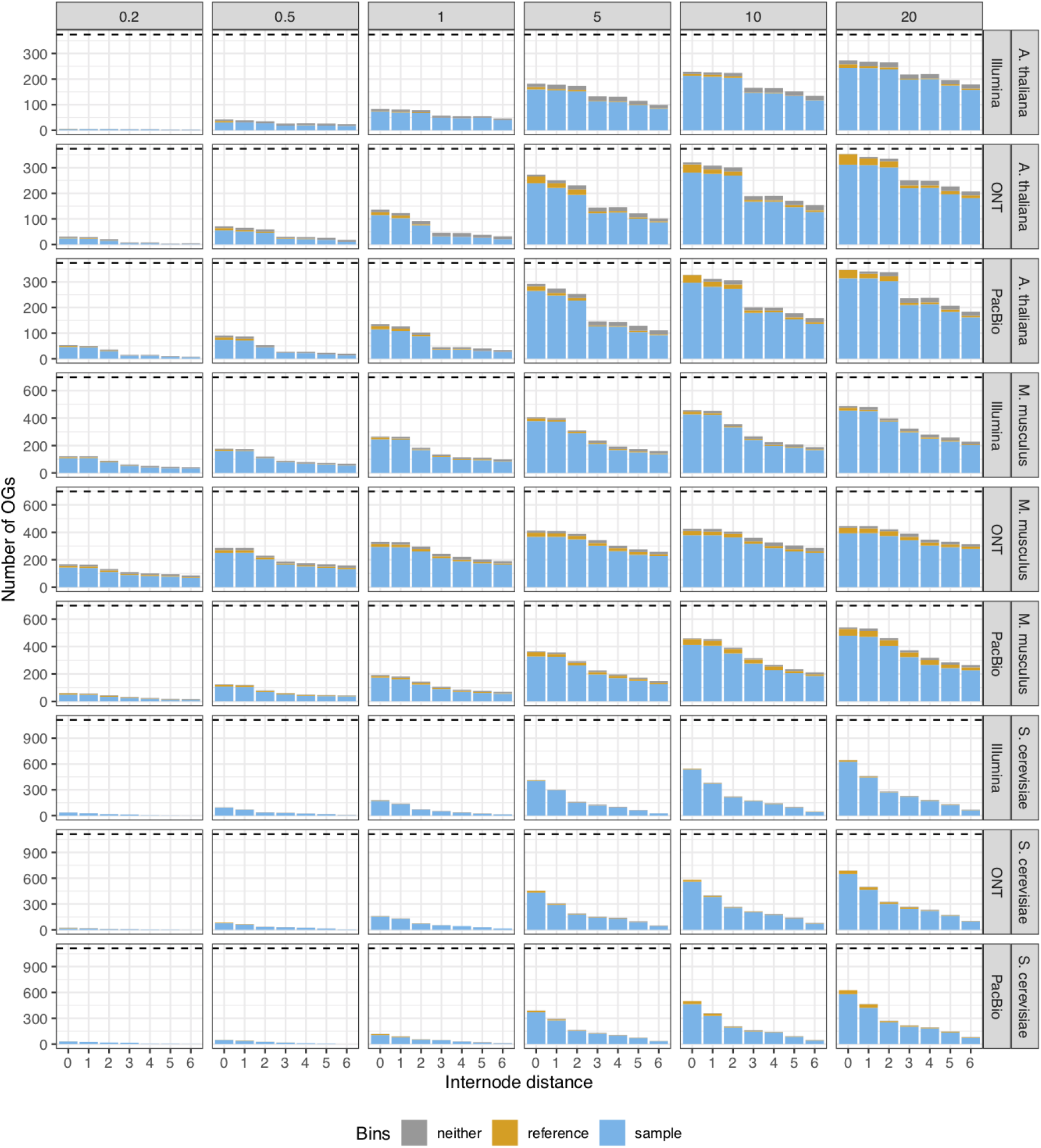
Binning of top BlastP results of r2t-sequence of selected species against their original OG (including the removed species) in either being most similar to its assembled counterpart (blue), to its reference used for reconstruction (yellow) or to any other sequence (grey). Results show that Read2Tree if reconstructing a sequence in most cases reconstructs a sequence that shows highest similarity to its assembled counterpart although this sequence was not present in the reference dataset when running Read2Tree.

**Supplementary Figure 4.**
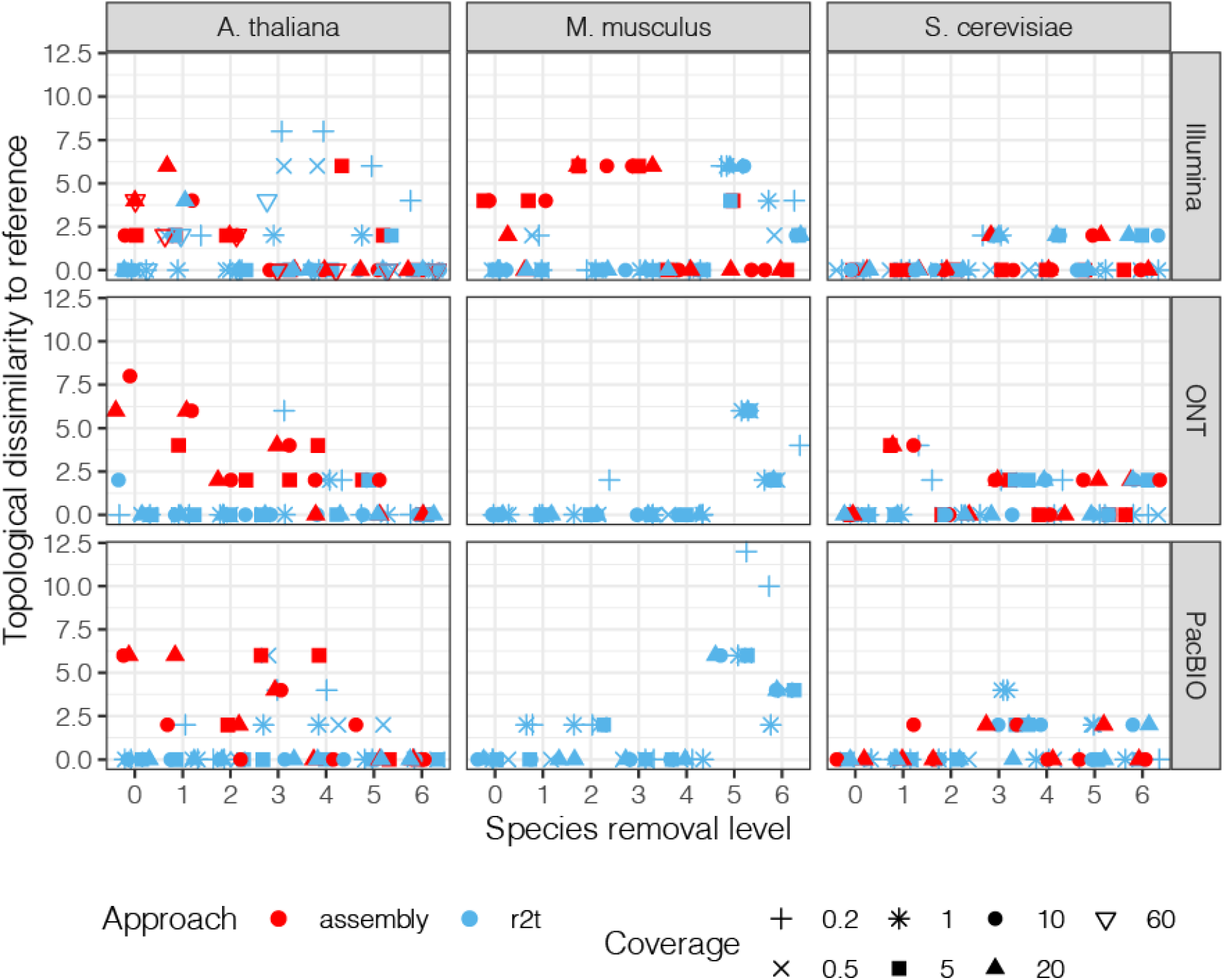
Comparison of Robinson Foulds tree distance of Read2Tree against reference tree and assembly obtained tree against reference tree. Read2Tree shows similar performance across technologies, coverage levels and distance to the closest remaining ancestor.

**Supplementary Figure 5.**
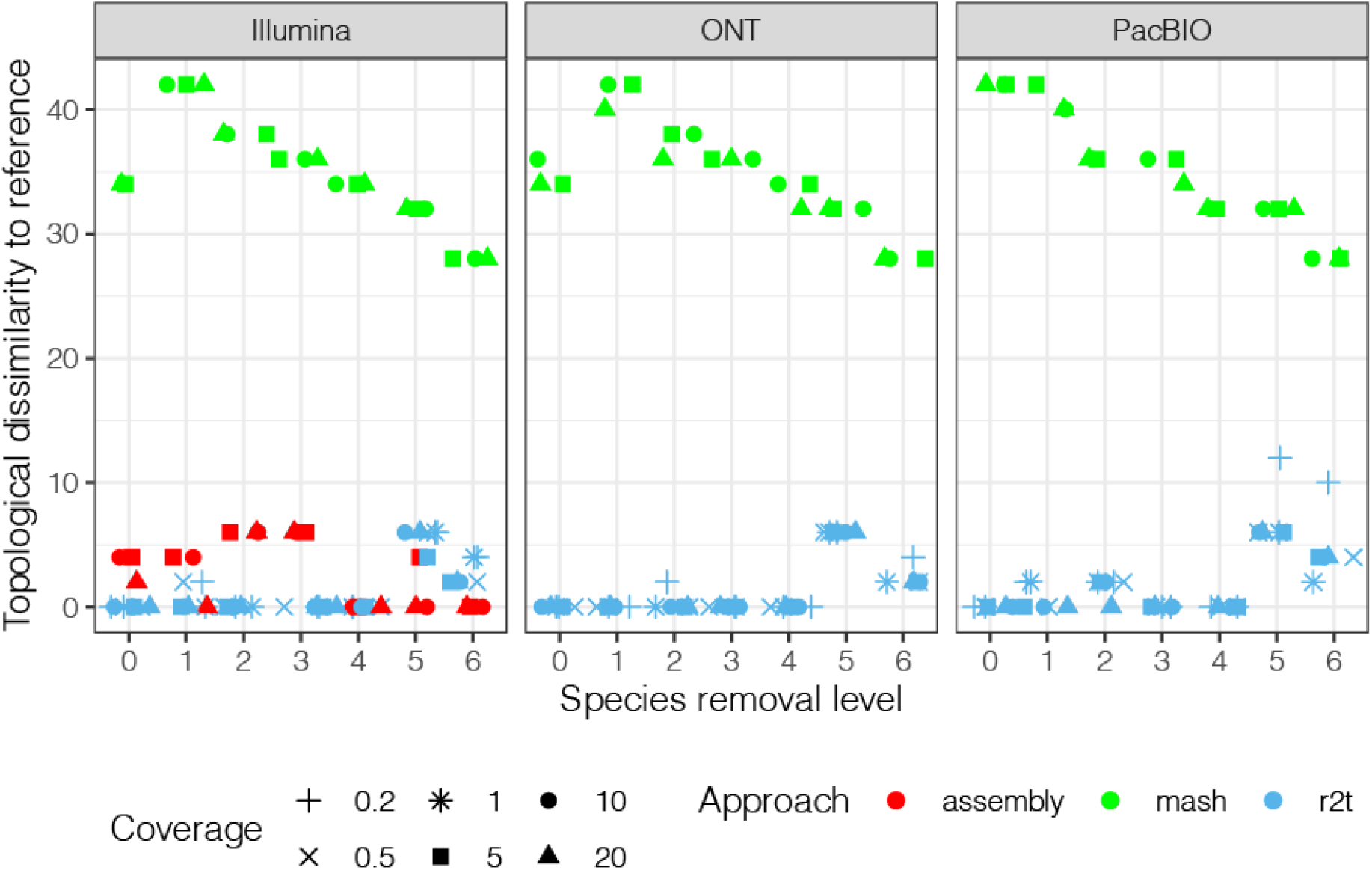
Topological dissimilarity (Robinson-Foulds distance) in comparison to reference trees for *M. musculus*. For the mash trees we have downloaded the genome assembly of species from NCBI Assembly. The Mash sketch with the size of 10m was used to create the k-mer sketch (k=21 as default) which was followed by mash distance to calculate the distances between genomes. Finally, RapidNJ was used on the distance matrix to infer the species tree. All trees are provided in Newick format in **Supplementary File 2.**

**Supplementary Figure 6.**
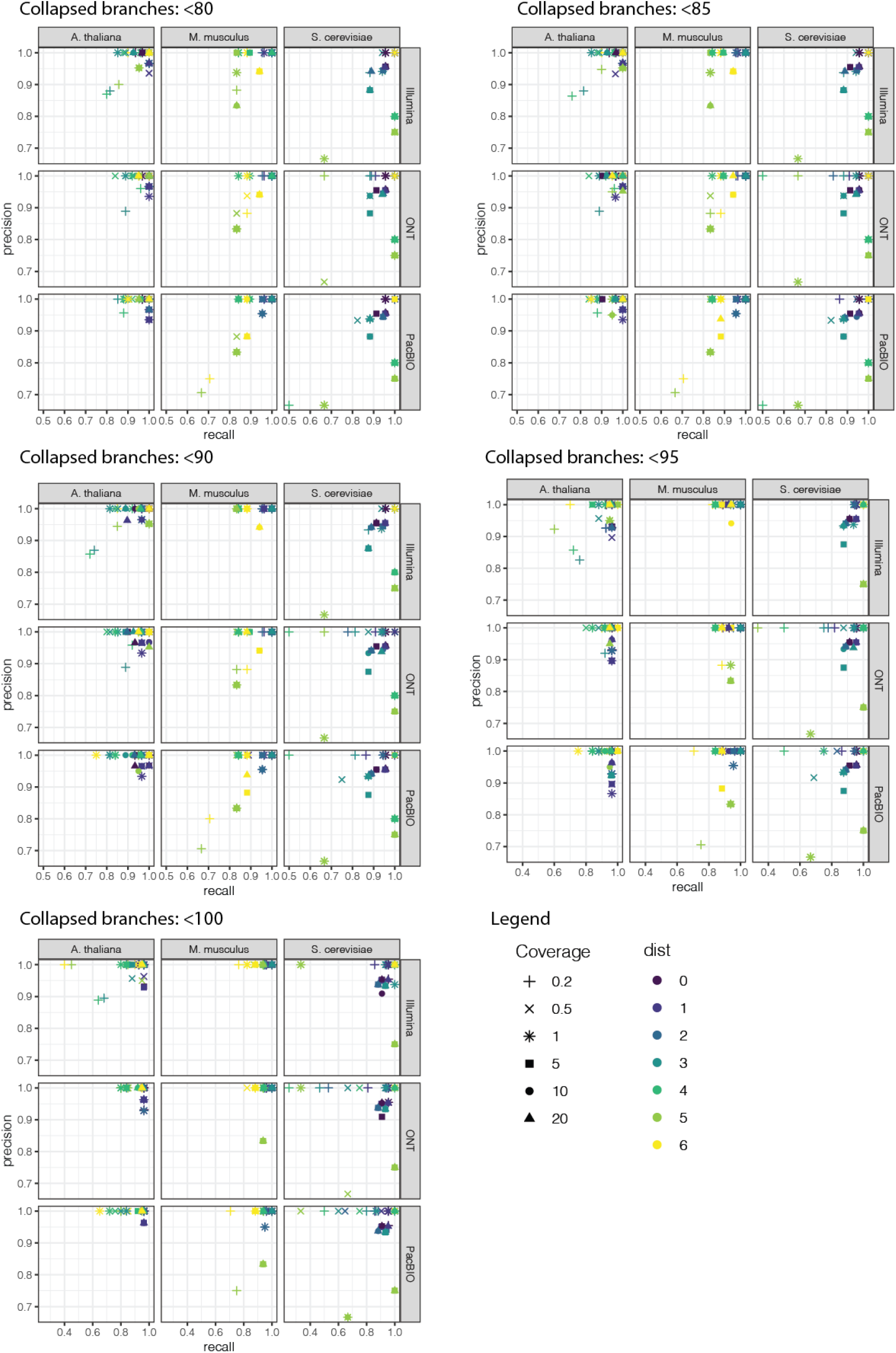
Precision recall plots for different levels of bootstraps for which branches were removed. Precision and recall increase for higher levels of bootstrap thresholds.

**Supplementary Figure 7.**
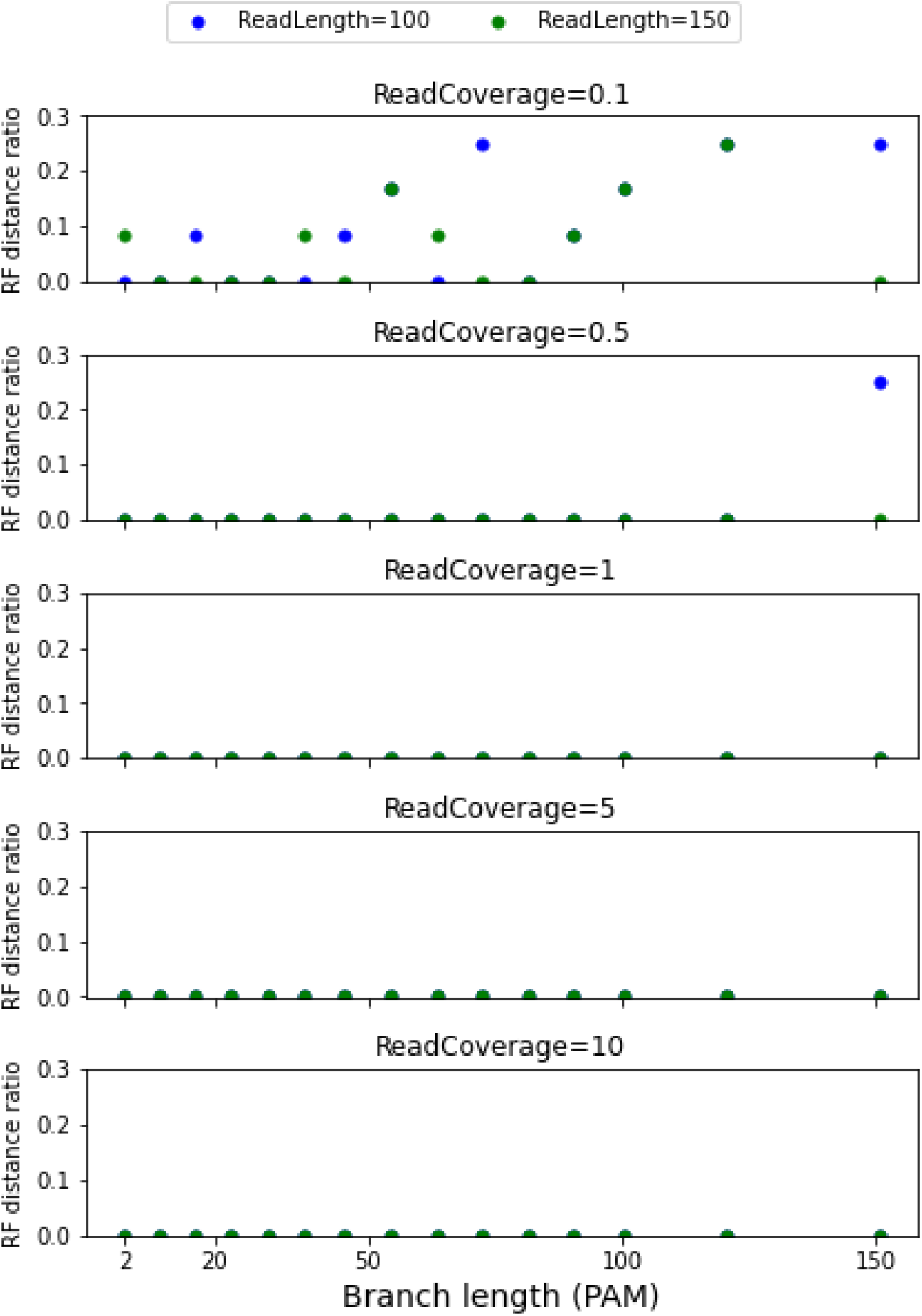
Comparison of Robinson Foulds tree distance of Read2Tree against true tree using simulated dataset. The design includes a fixed topology for species tree with 15 species using the ALF package. We varied the branch length leading to one of the species (species of interest) between 2 PAM and 150 PAM. For each run, we infer afterwards the OMA Groups (excluding the species of interest). Then using art_illumina, we generated DNA sequencing reads (paired-end) with length of 100 and 150 and coverage of 0.1 to 10. Next, for each case, we ran the read2tree package to infer the phylogeny. Finally, we calculated the Robinson–Foulds metric between inferred species tree and the true one (output of ALF). As one can see from the figure, only in 0.1x coverage inferring the true tree is challenging, otherwise in almost all the cases the tree is inferred perfectly.

**Supplementary Figure 8.**
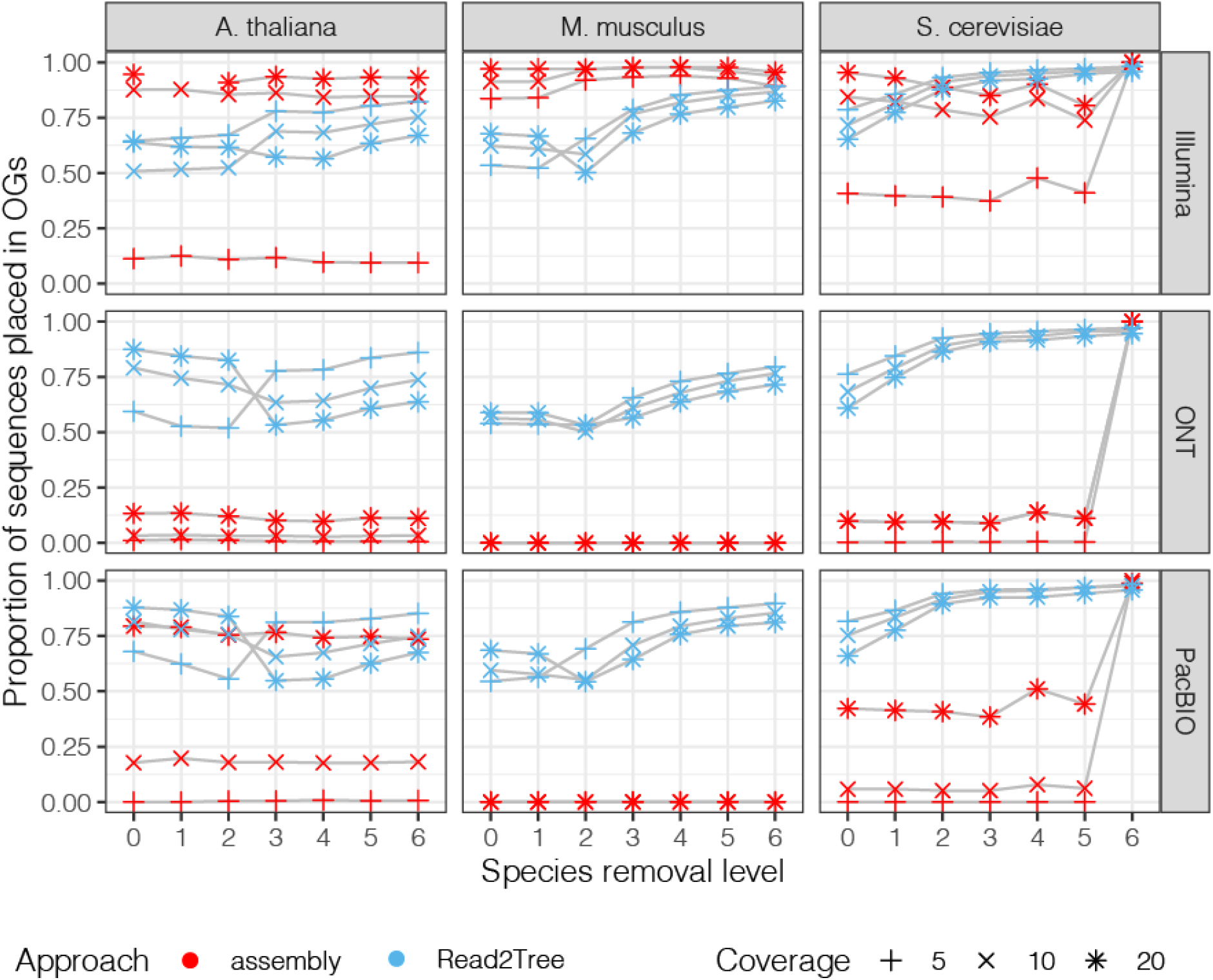
Proportions of sequences placed into the total number of OGs when selecting OGs with at least 80% taxa.

**Supplementary Figure 9.**
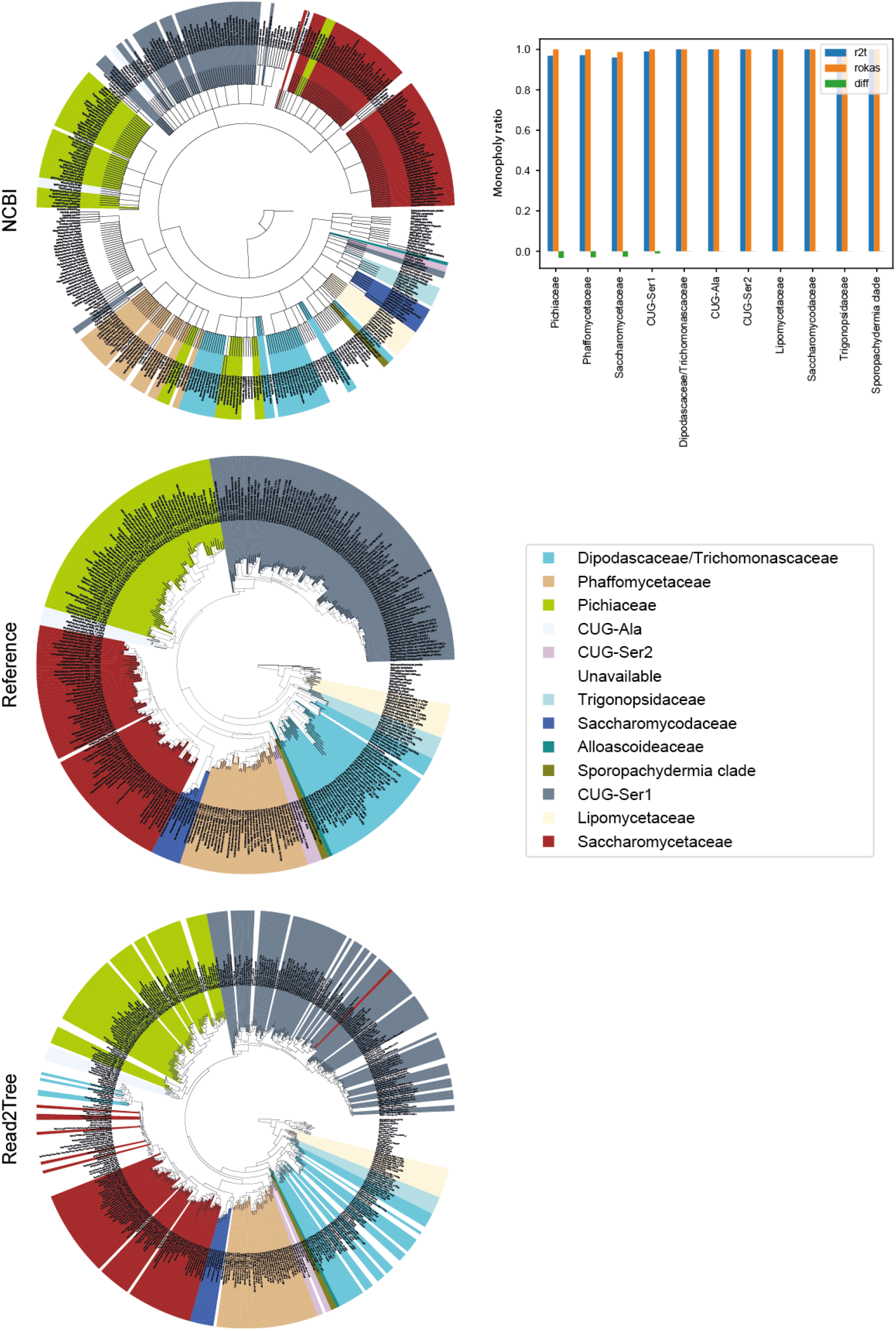
Comparison of Trees by classification based on (Shen et al., 2018) at order level. Left NCBI, reference and Read2Tree inferred trees. Right Monophyly score computed for Shen *et al*. classification. Read2Tree is nearly as precise and the standard pipeline in recapitulating the right monophyletic groupings.

**Supplementary Figure 10.**
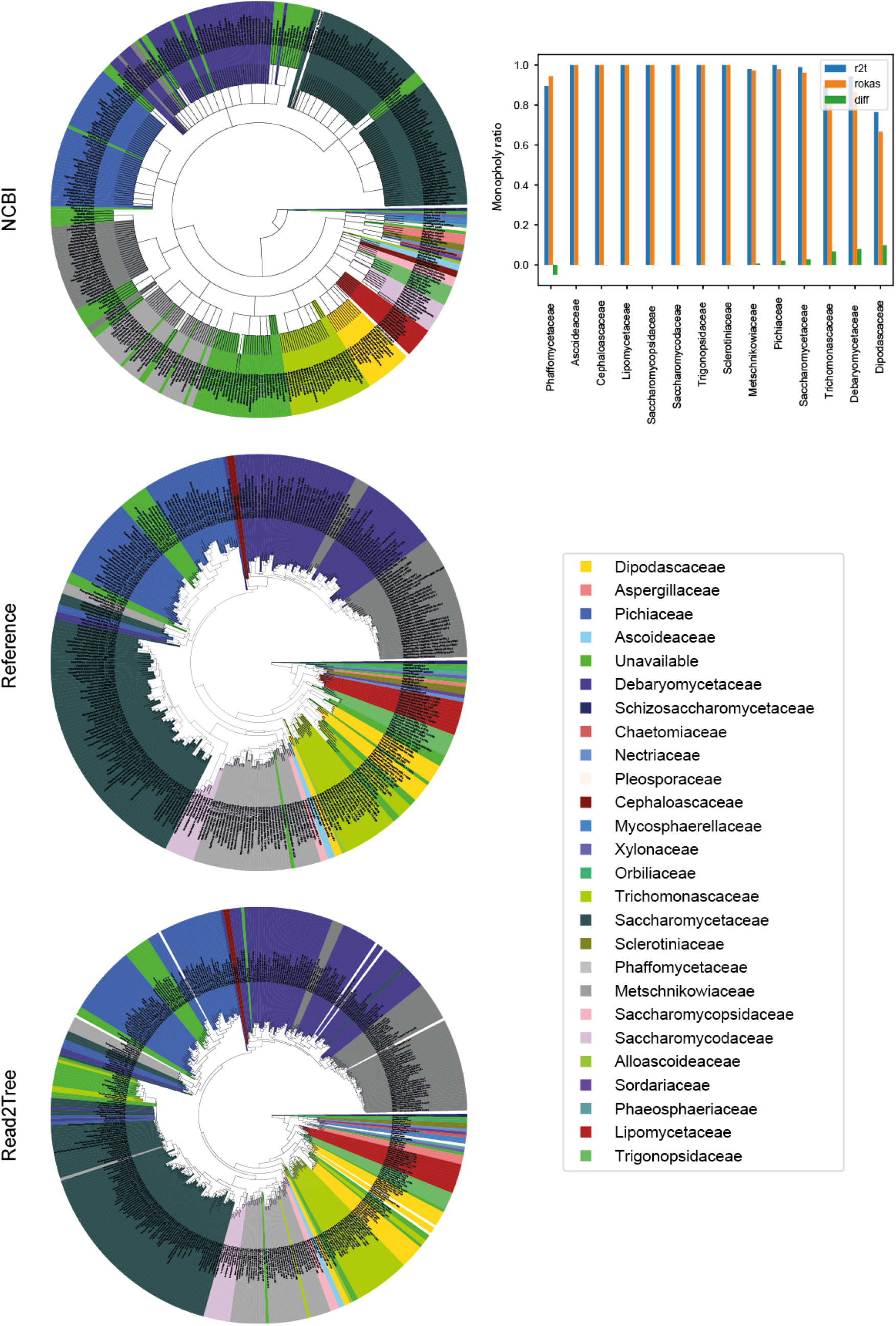
Comparison of Trees by classification based on (Shen et al., 2018) at family level. Left NCBI, reference and Read2Tree inferred trees. Right Monophyly score computed for Shen *et al*. classification. Read2Tree is nearly as precise and the standard pipeline in recapitulating the right monophyletic groupings. Similar comparison at order level is provided in Supplementary Figure 9.

**Supplementary Figure 11.**
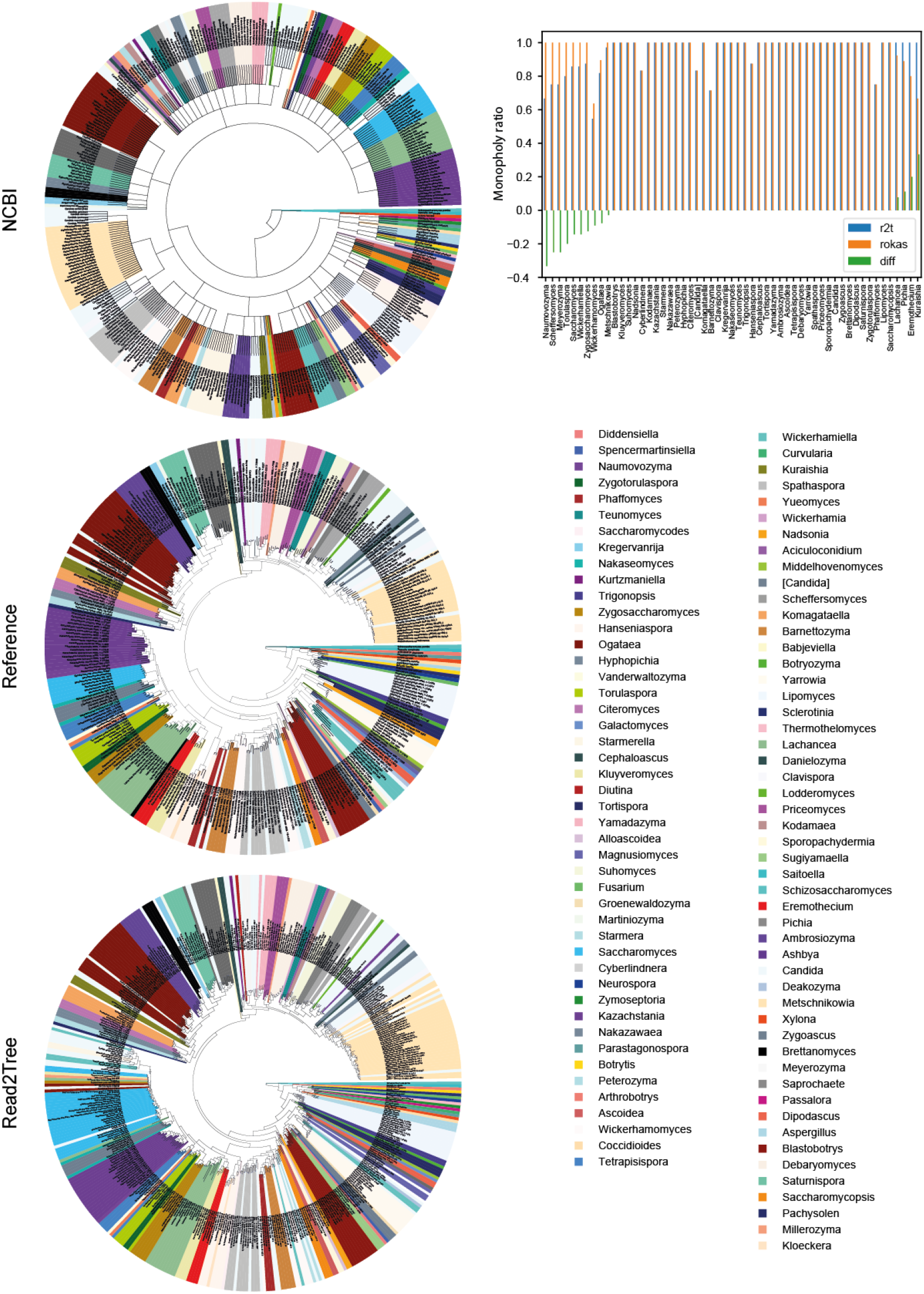
Comparison of Trees by classification based on (Shen et al., 2018) at genus level. Left NCBI, reference and Read2Tree inferred trees. Right Monophyly score computed for Shen *et al*. classification. Read2Tree is nearly as precise and the standard pipeline in recapitulating the right monophyletic groupings.

**Supplementary Figure 12.**
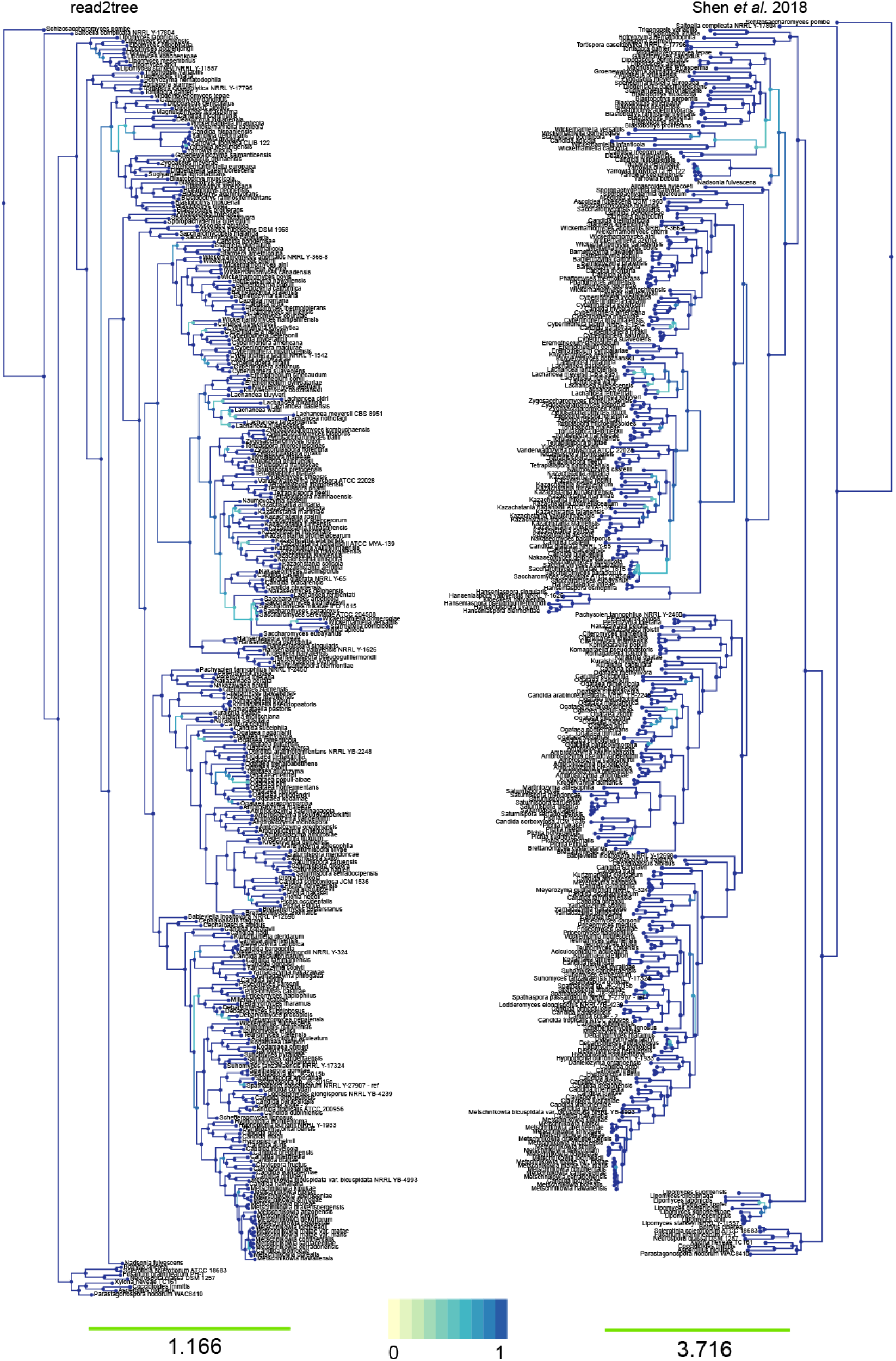
Side by side comparison of read2tree using phylo.io. In dark blue are all the branches that are equal between the two trees using the best corresponding node algorithm.

**Supplementary Figure 13 .**
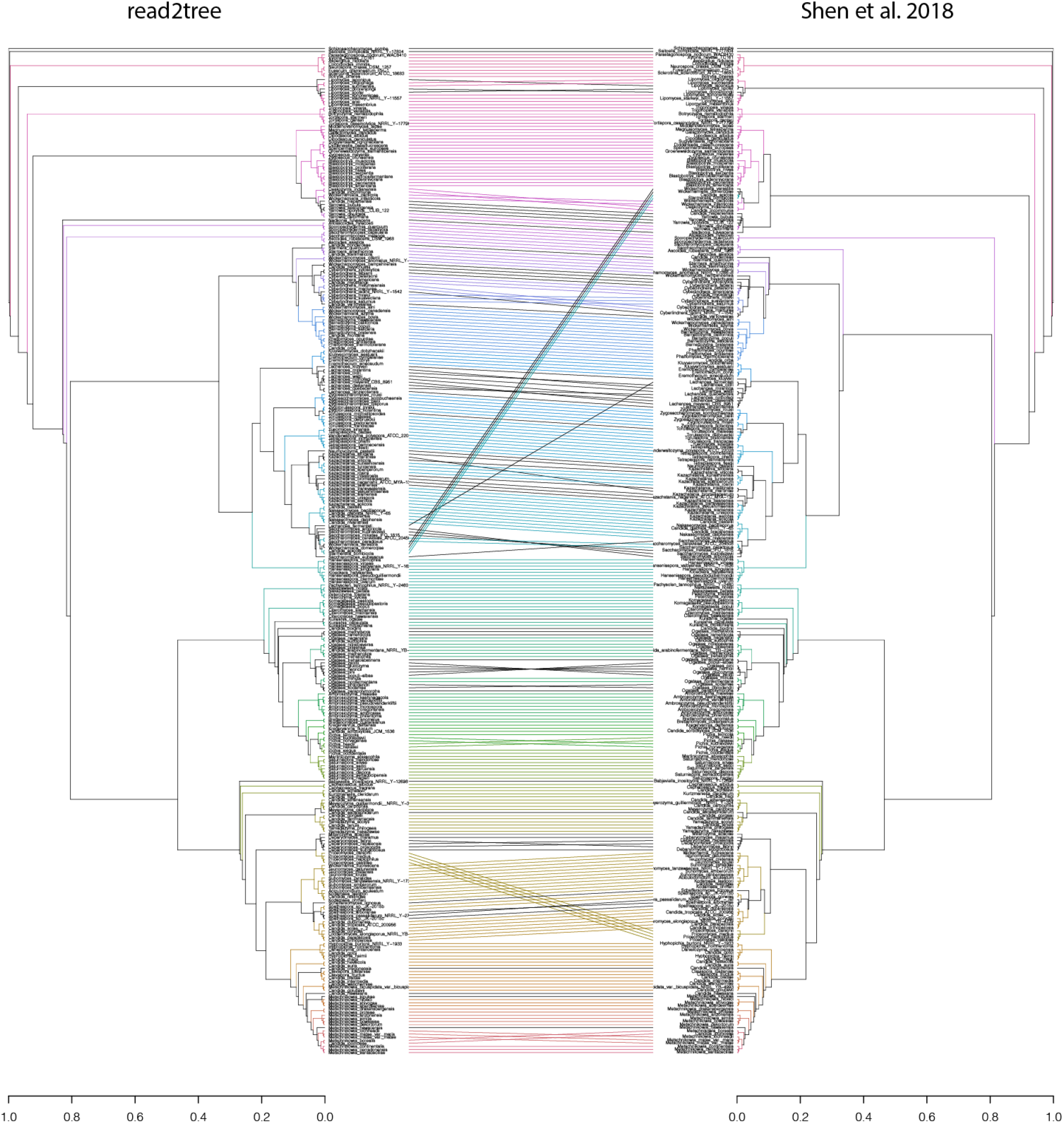
Tanglegram of yeast tree comparison highlighting differences and similarities in the tree. Entanglement coefficient was 0.0096. Tanglegram was produced using the dendextend R library.

**Supplementary Figure 14 .**
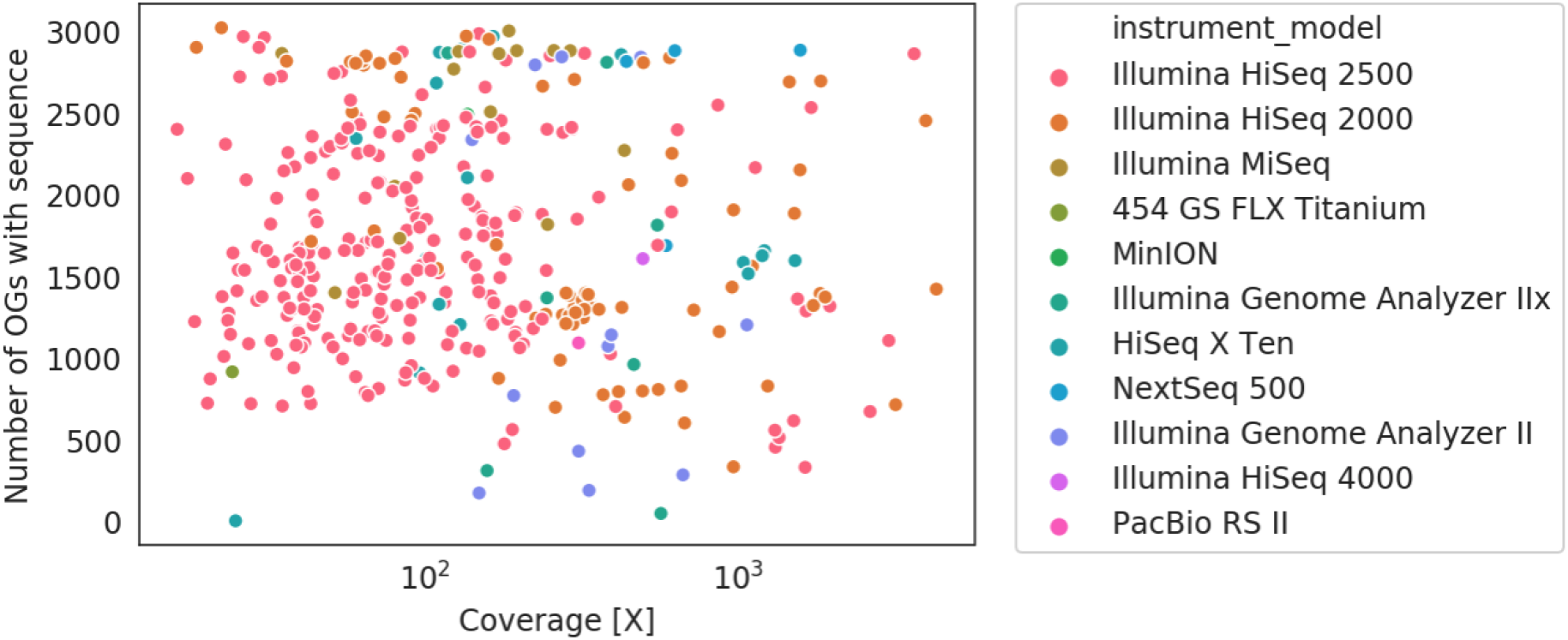
Coverage to the number of obtained sequence relationships. Even for low coverage we see a large number of obtained sequences. No clear relationship is present between input coverage and number of sequences placed in OGs. In purple at 100X we display the number of sequences present in all ~3000 OGs for the reference species.

**Supplementary Figure 15.**
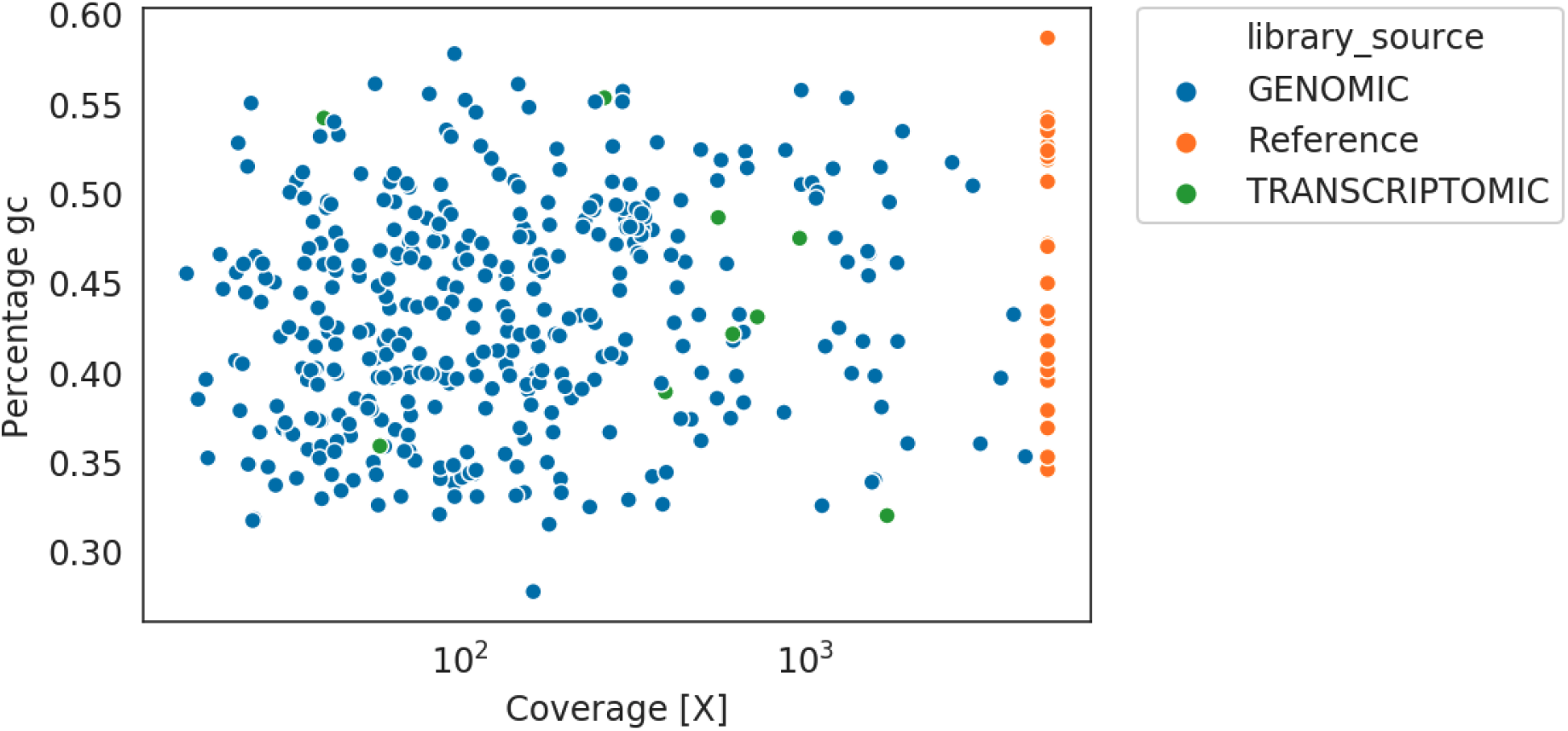
Average percentage GC content in obtained sequences for yeast tree obtained using Read2Tree in comparison to given references. In most cases the reconstructed sequences are within the range of GC content as present in the reference sequences. References sequence artificially set to 5000X coverage for display purposes.

**Supplementary Figure 16.**
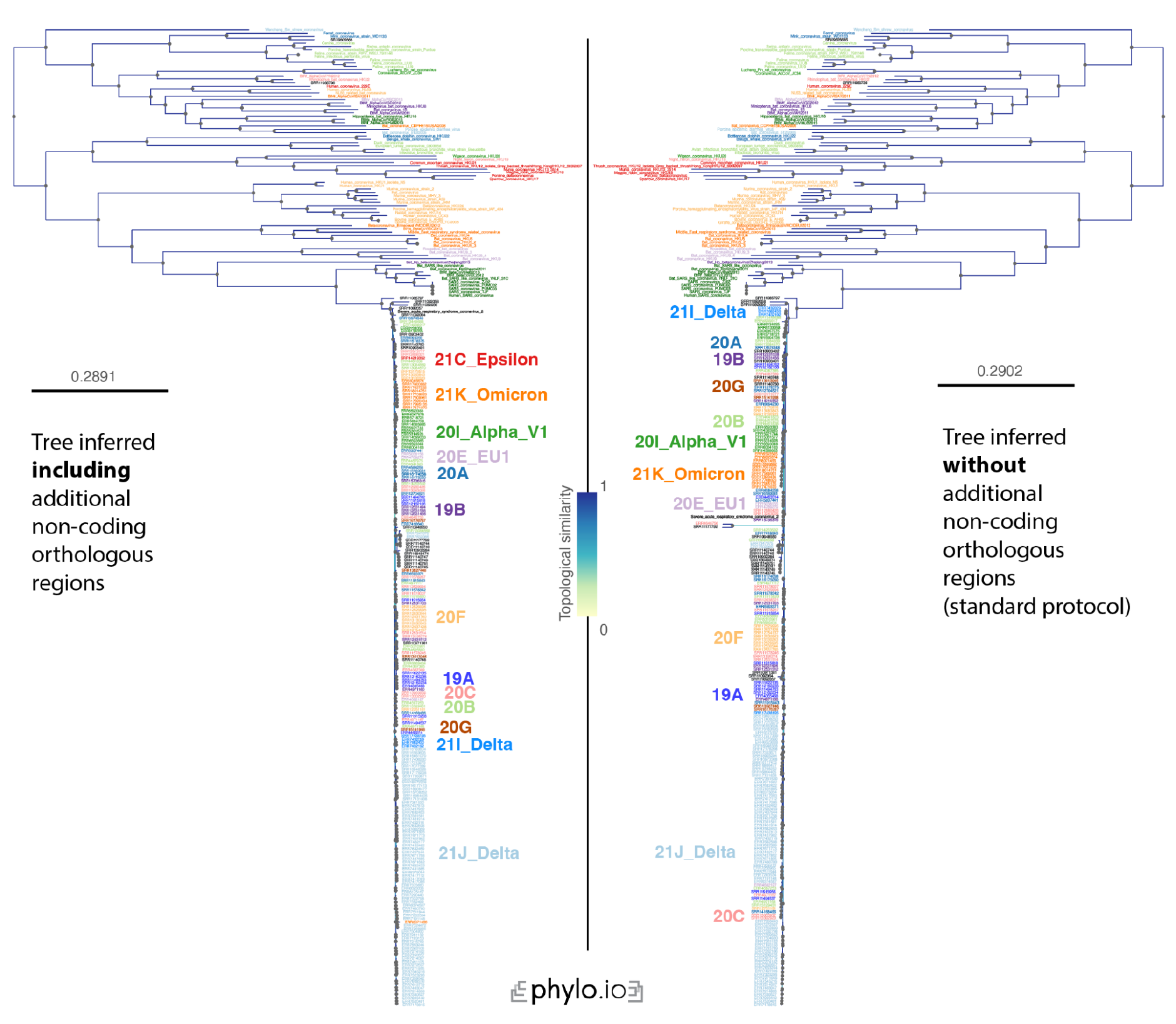
Comparison of Read2Tree-reconstructed Coronaviridae trees with (left) and without (right) additional non-coding reference regions (see Methods). The backbone is 100% consistent, while the SARS-CoV-2 part of the tree shows a bit more variation. Nevertheless, the colour clustering in both trees shows that Read2Tree can be used to reliably classify samples according to the fine-grained Nextstrain (Hadfield et al., 2018) classification. Best performance is obtained when including additional non-coding reference markers as described in the *Methods* section.

**Supplementary Figure 17.**
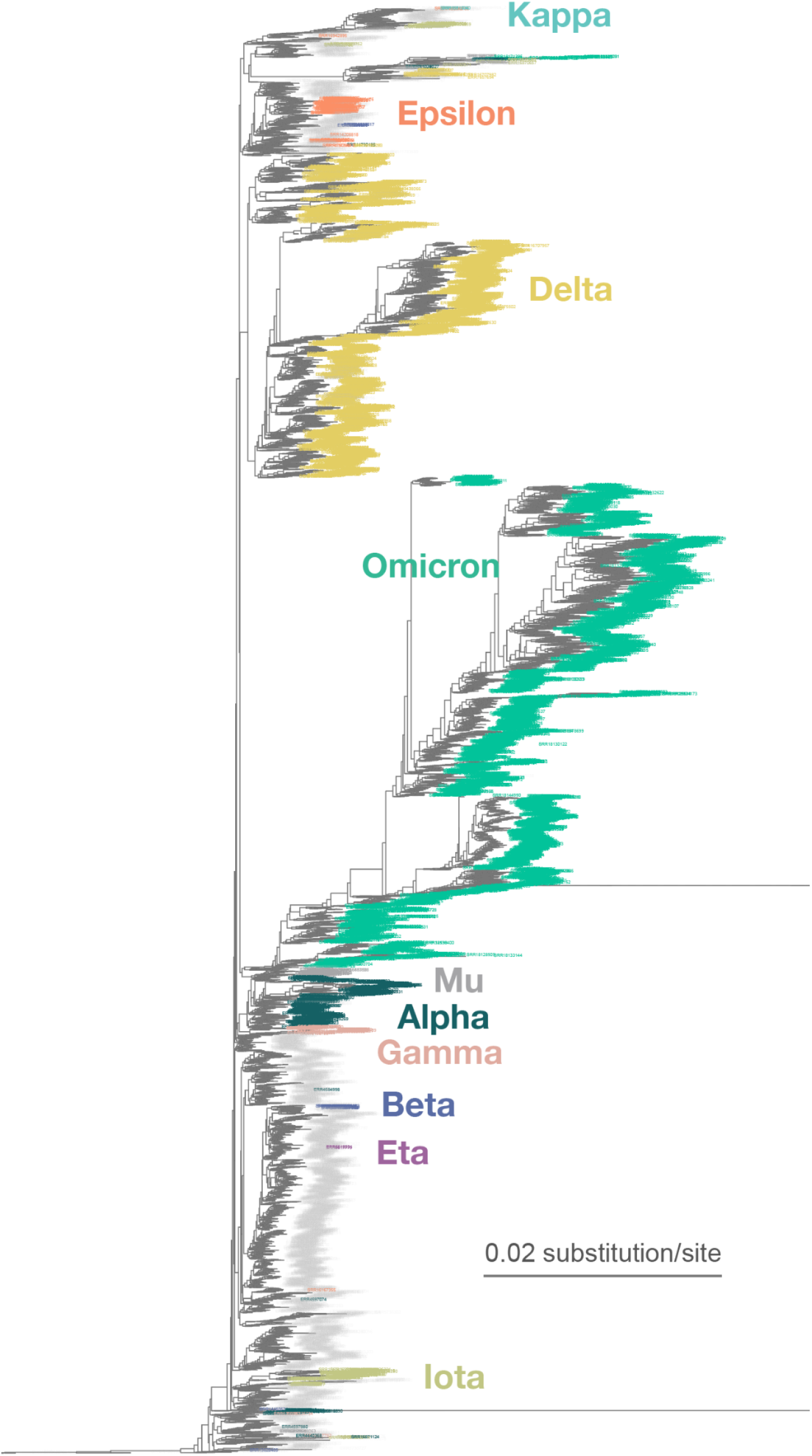
Zoomed-in display of a tree inferred using Read2Tree on 10,283 samples whole genome SARS-CoV-2 samples. Classification in colour was obtained from https://harvestvariants.info/ (see Methods), where grey leaves are unclassified according to the CDC label. The colour clustering shows that the Read2Tree-based tree recovers consistent classification.

**Supplementary Figure 18.**
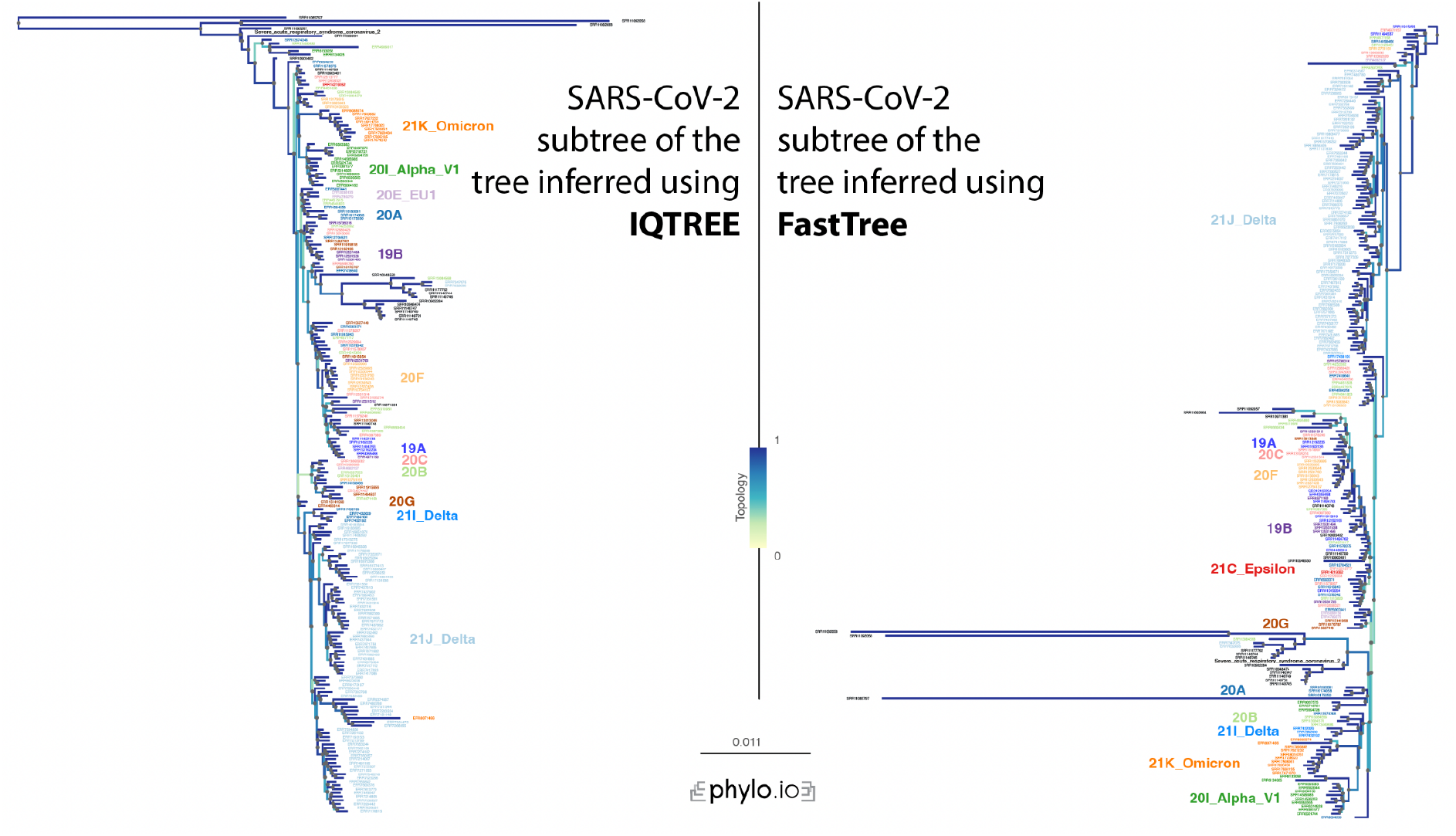
Comparison of IQTree and FastTree for Coronaviridae. Classification can be done quite reliably with both tree inference methods, but the rooting appears to be incorrect in the FastTree-reconstructed tree. The two trees are provided in the Supplementary File 2 in Newick format.

