## Supplementary material for "Read2Tree: scalable and accurate phylogenetic trees from raw reads": Previous supplement

### Supplementary Methods

#### Detailed description of Read2Tree

**Data Inputs**
Read2Tree requires two sets of inputs: 1. A set of reads for a species of interest and 2. Reference orthologous groups.

**Description of read2tree**
In a first step Read2Tree pulls the DNA sequences for the obtained OGs from the oma database using the available API and separates the OGs into individual folders. Then it collects for each species the DNA sequences and produces one file per species with the relevant sequences. These files are then used for the mapping. To allow for parallelization mapping can be performed on all species sequentially or set individually allowing to span multiple jobs per reference species on an HPC. Mapping of sequences is performed using NextGenMap (Sedlazeck et al., 2013) for short reads and NextGenMap-LR for long reads (Sedlazeck et al., 2018). Mapped reads are then post processed using samtools to extract the bases in regards to the sequences. Once the bases for a given reference sequence are extracted the consensus sequence is built based on the majority base at a position requiring at least 3 reads to be present. The reconstructed sequences are then placed back to the original OG and alignment. All alignments of OGs are then concatenated and result in the final supermatrix that serves as input for any tree inference method.

### Supplementary Figures

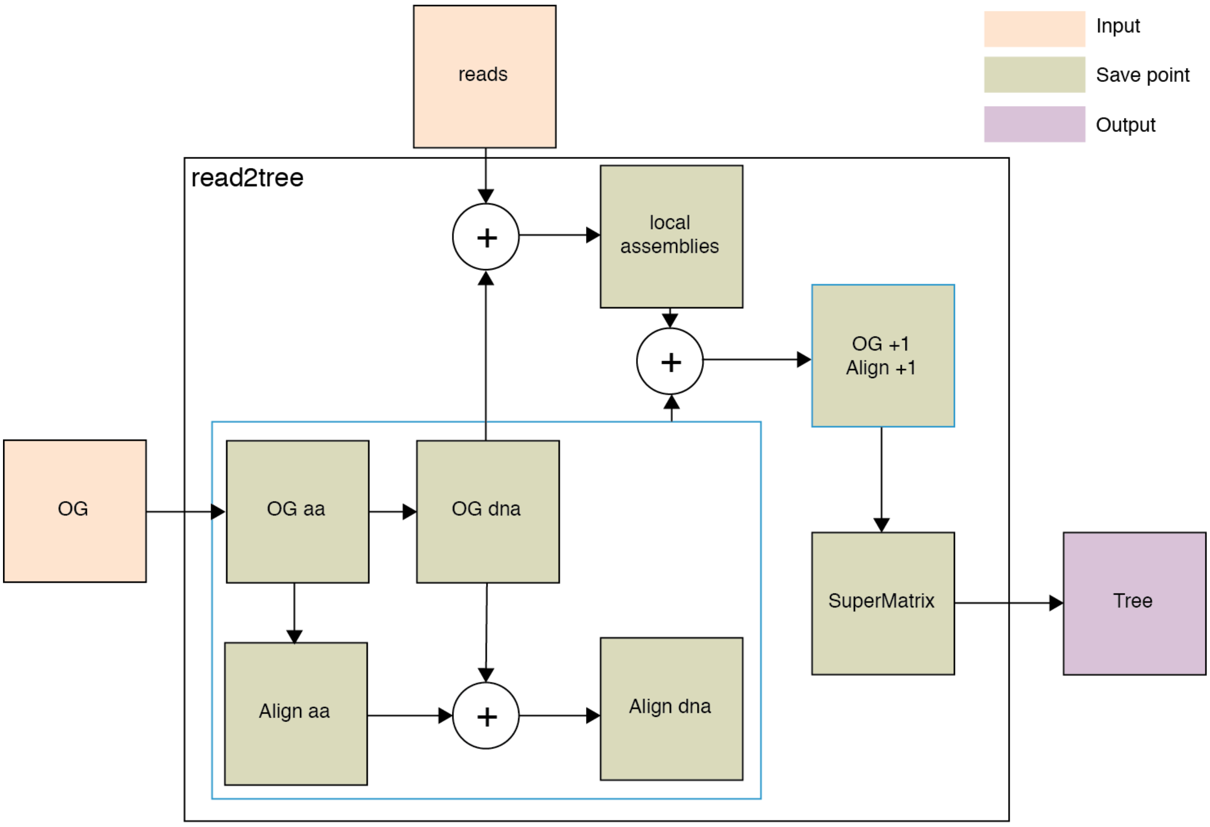


**Supplementary Figure 1.** Graphical representation of pipeline. All boxes in green are stored by Read2Tree. Inputs are reads and a set of reference orthologous groups that can be selected from over 2000 species from the OMA database. Local assemblies here as reconstructed sequences using the bases placed against the reference.

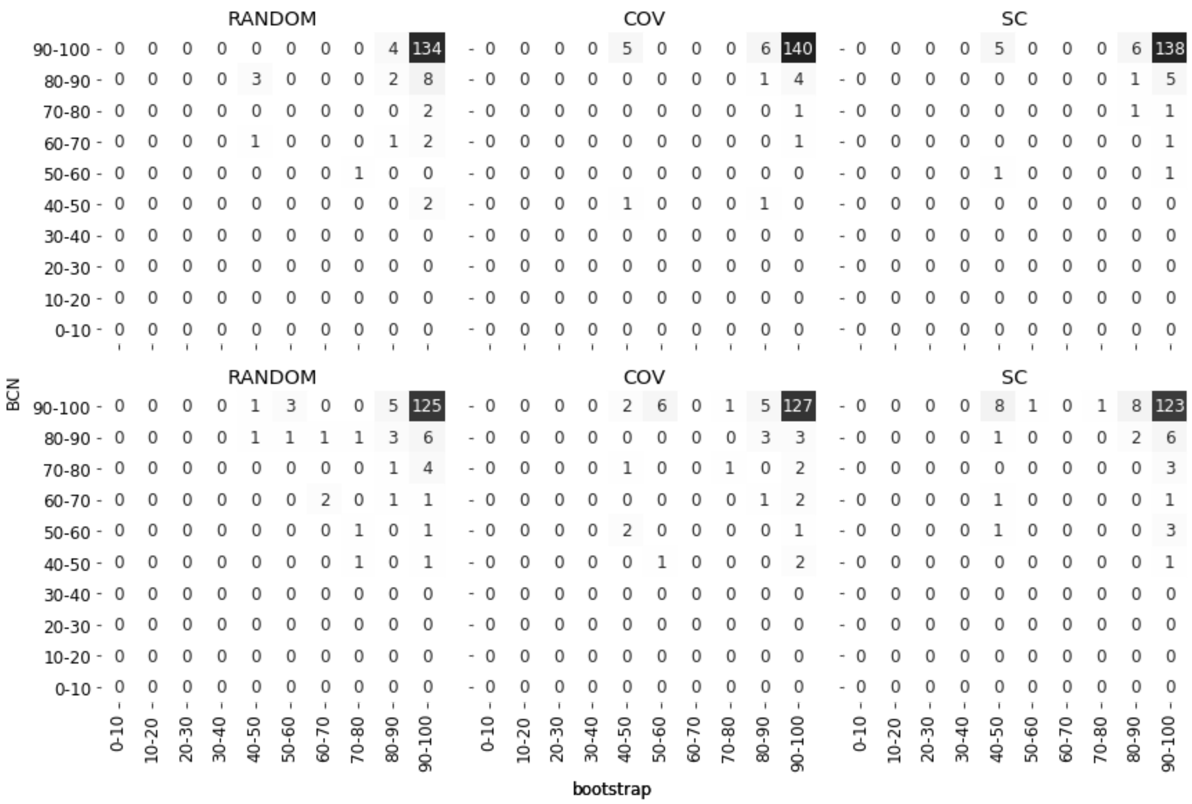


**Supplementary Figure 2.** Comparison of different gene selection methods in 20 times 8out/8in test show differences in relation between bootstrap on nodes of mapped species and its jaccard similarity with reference tree (Best Corresponding Node BCN) value. Top row shows values obtained using simulated reads and bottom row shows values obtained using real reads. In both cases we see that using coverage as a selection method shows the highest number of nodes where mapped species has high bootstrap and high BCN value.

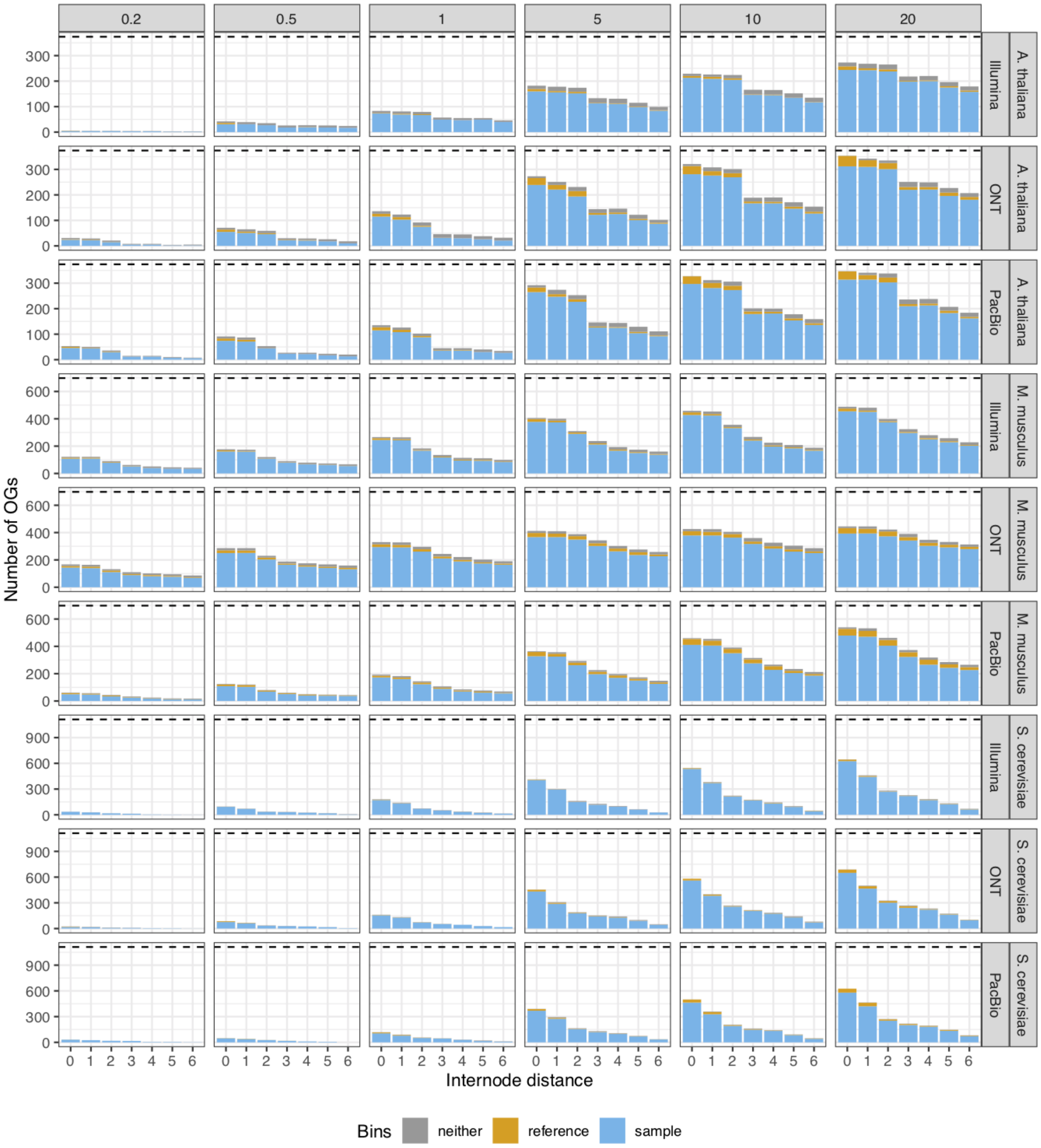


**Supplementary Figure 3.** Binning of top blastp results of r2t-sequence of selected species against their original OG (including the removed species) in either being most similar to its assembled counterpart (blue), to its reference used for reconstruction (yellow) or to any other sequence (grey). Results show that Read2Tree if reconstructing a sequence in most cases reconstructs a sequence that shows highest similarity to its assembled counterpart although this sequence was not present in the reference dataset when running Read2Tree.

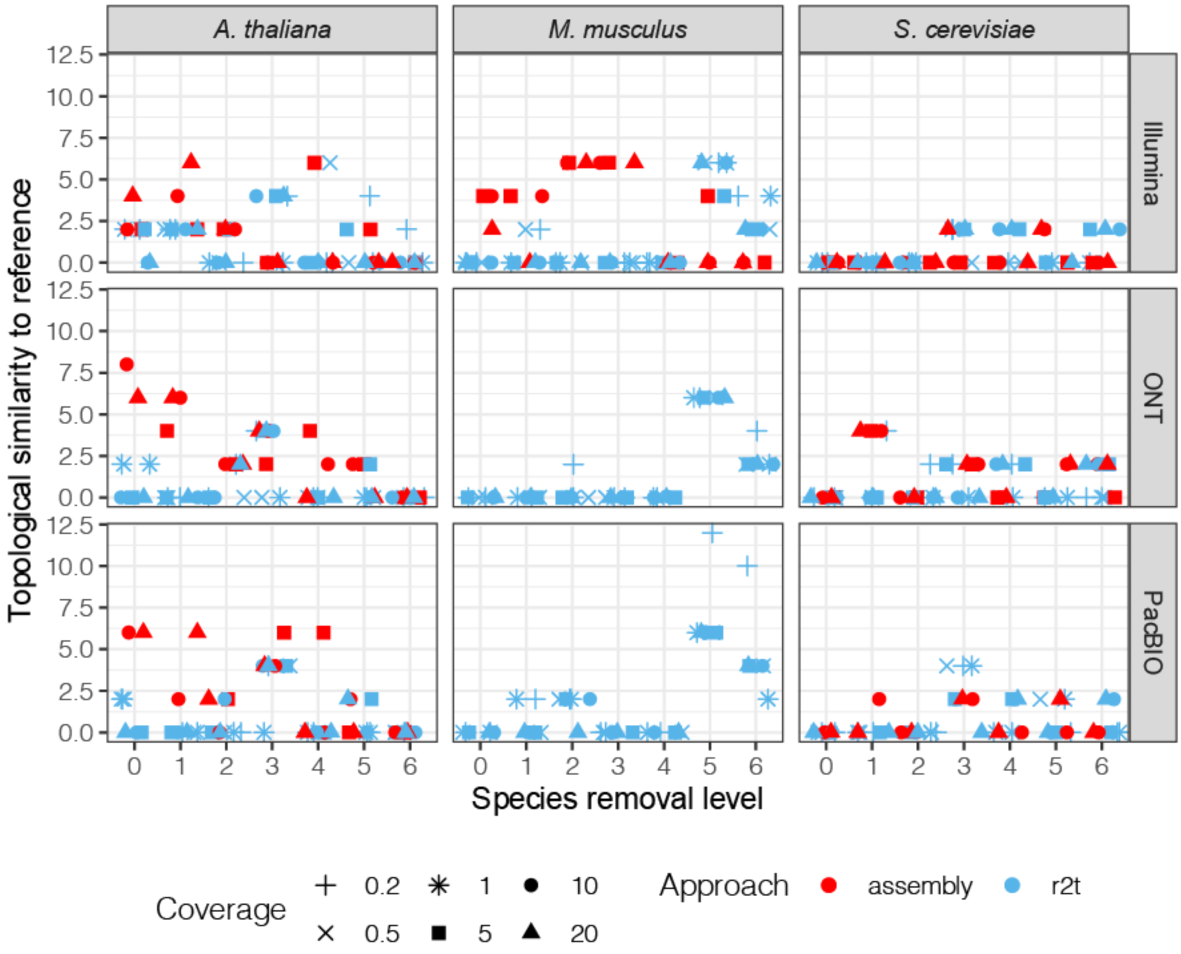


**Supplementary Figure 4.** Comparison of robinson foulds tree distance of Read2Tree against reference tree and assembly obtained tree against reference tree. Read2Tree shows similar performance across technologies, coverage levels and distance to the closest remaining ancestor.

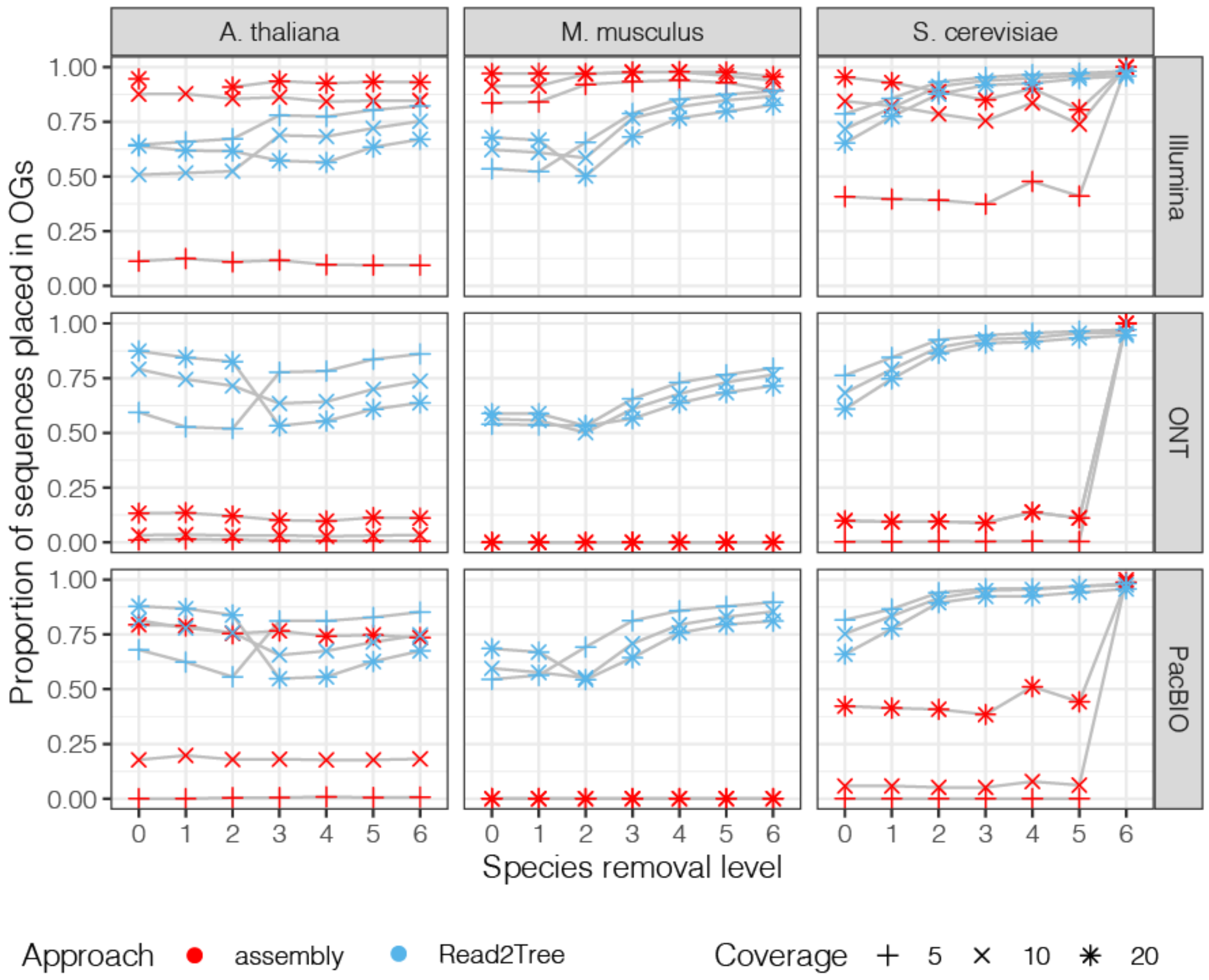


**Supplementary Figure 5.** Proportions of sequences placed into the total number of OGs when selecting OGs with at least 80% taxa.

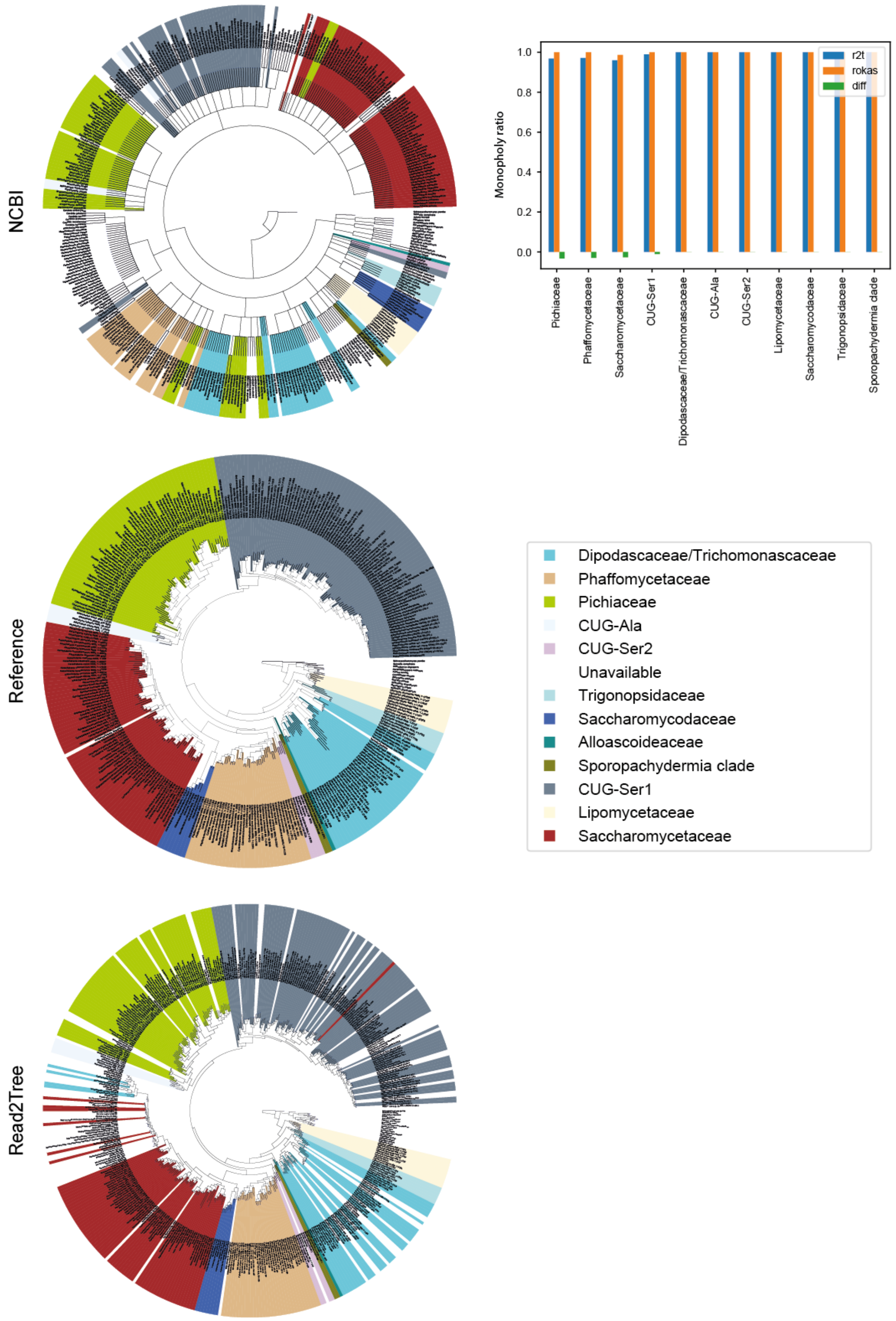


**Supplementary Figure 6.** Comparison of Trees by classification based on (Shen et al., 2018). Left NCBI, reference and Read2Tree inferred trees. Right Monophyly score computed for Shen *et al.* classification. Read2Tree is nearly as precise and the standard pipeline in recapitulating the right monophyletic groupings.

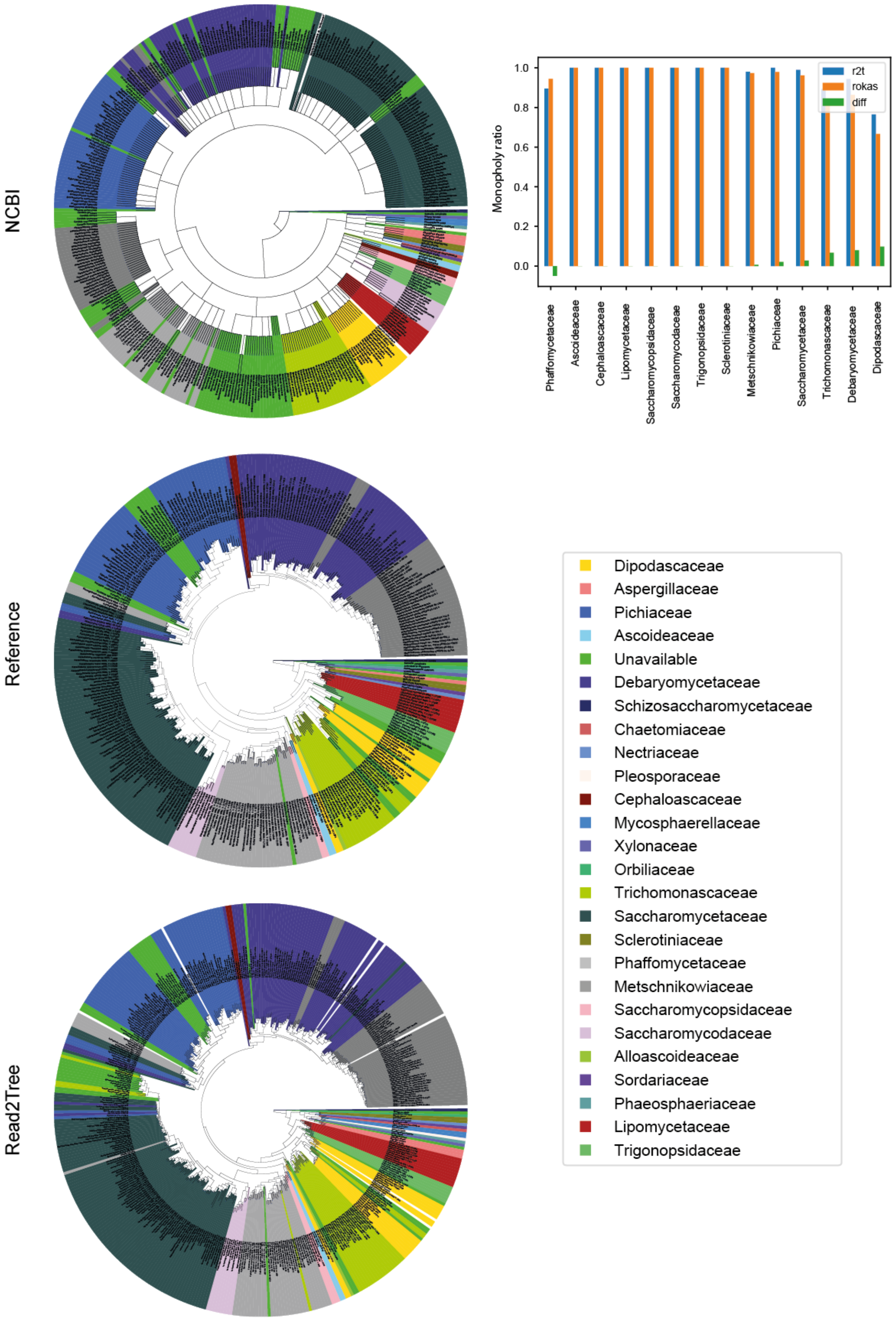


**Supplementary Figure 7.** Comparison of Trees by classification based on (Shen et al., 2018). Left NCBI, reference and Read2Tree inferred trees. Right Monophyly score computed for Shen *et al.* classification. Read2Tree is nearly as precise and the standard pipeline in recapitulating the right monophyletic groupings.

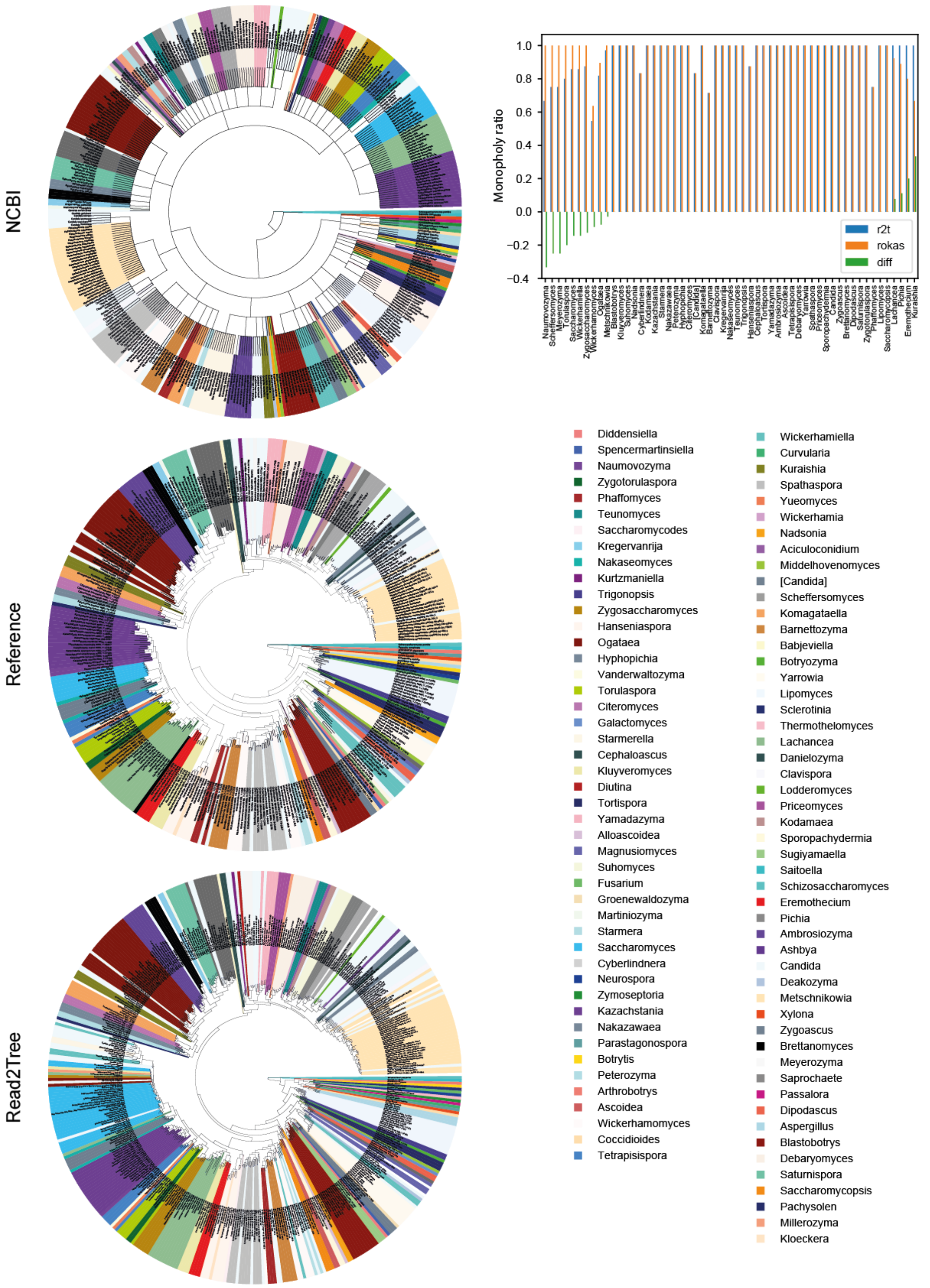


**Supplementary Figure 8.** Comparison of Trees by classification based on (Shen et al., 2018).Left NCBI, reference and Read2Tree inferred trees. Right Monophyly score computed for Shen *et al.* classification. Read2Tree is nearly as precise and the standard pipeline in recapitulating the right monophyletic groupings.

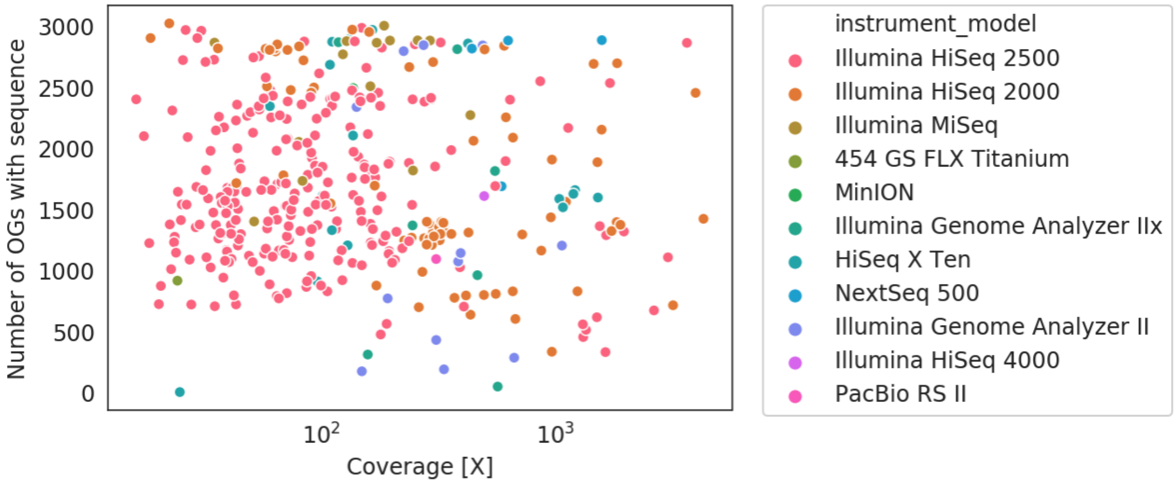


**Supplementary Figure 9.** Coverage to the number of obtained sequence relationships. Even for low coverage we see a large number of obtained sequences. No clear relationship is present between input coverage and number of sequences placed in OGs. In purple at 100X we display the number of sequences present in all ~3000 OGs for the reference species.

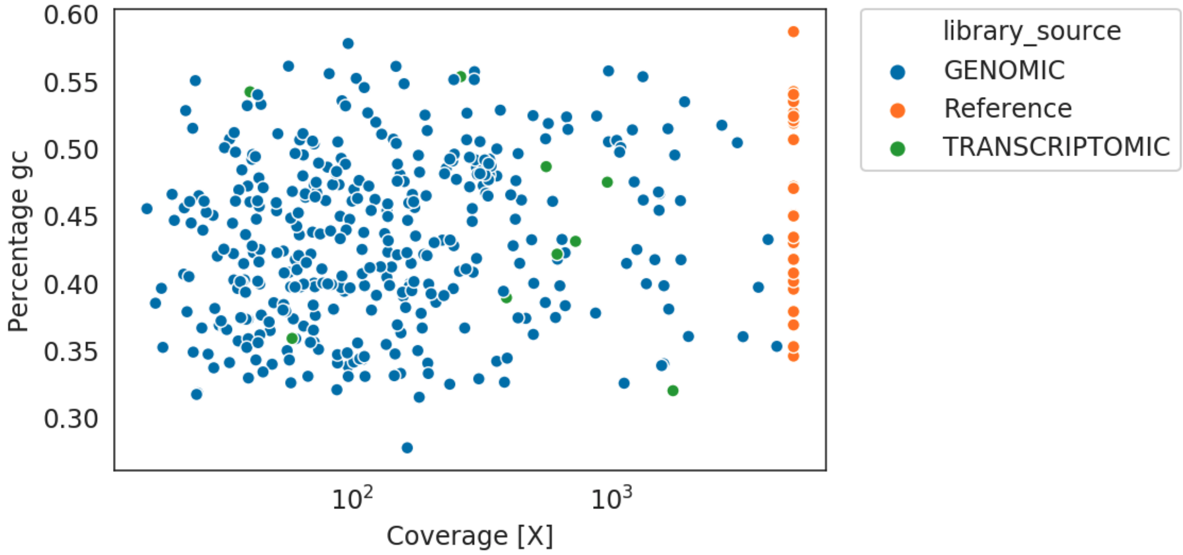


**Supplementary Figure 10.** Average percentage GC content in obtained sequences for yeast tree obtained using Read2Tree in comparison to given references. In most cases the reconstructed sequences are within the range of GC content as present in the reference sequences. References sequence artificially set to 5000X coverage for display purposes.

### Tables

#### Reads used for species placement test

| Species | Machine | BasePairs | Source | Accession |
|---|---|---|---|---|
| *A. thaliana* | PacBio | 7.0 Gbp | DNA | ERR2173371 |
| *A. thaliana* | MinION | 3.4 Gbp | DNA | ERR2173373 |
| *A. thaliana* | Illumina MiSeq | 8.4 Gbp | DNA | ERR2173372 |
| *M. musculus* | Illumina HiSeq | 5.1 Gbp | mRNA | SRR5171076 |
| *M. musculus* | PacBio | 2.716 Gbp | mRNA | SRR5314792<br>SRR5314793<br>SRR5314794<br>SRR5314795<br>SRR5314796<br>SRR5314797<br>SRR5314798 |
| *M. musculus* | MinION | 0.7892 Gbp | mRNA | SRR4048177<br>SRR4048178<br>SRR4048179<br>SRR4095033<br>SRR4095035<br>SRR5286956<br>SRR5286957<br>SRR5286958<br>SRR5286959<br>SRR5286960<br>SRR5286961<br>SRR5286962<br>SRR5286963 |
| *S. cerevisiae* | Illumina HiSeq | 2.5 Gbp | DNA | SRR5892450 |
| *S. cerevisiae* | PacBio | 4.9 Gbp | DNA | SRR5989371 |
| *S. cerevisiae* | MinION | 1.129 Gbp | DNA | SRR5892449<br>SRR5924195 |

#### Reference datasets

| **Species** | **#Ref** | **#OGs** | **#min species OG** | **Outgroup** | **#OGs with Species** |
|---|---|---|---|---|---|
| MOUSE | 30 | 699 | 26 | *Amphimedon queenslandica* | 698 |
| YEAST | 26 | 1188 | 21 | *Monosiga brevicollis* | 1133 |
| ARATH | 35 | 372 | 32 | *Chlorella variabilis* | 369 |

#### Reads used for Figure 2 and 3

See additional file supplementary_file_1.csv

#### Reads used for Figure 4

See additional file supplementary_file_1.csv

#### Reference dataset for mapping of 435 budding yeast species

Reference dataset was obtained from the OMA database (Altenhoff et al., 2021) using marker gene export function for 31 species with minimum coverage threshold of 0.8.

See additional file yeast_reference.csv

Sedlazeck FJ, Rescheneder P, Smolka M, Fang H, Nattestad M, von Haeseler A, Schatz

MC. 2018. Accurate detection of complex structural variations using single-molecule sequencing. *Nat Methods* **15**:461–468. doi:10.1038/s41592-018-0001-7

Sedlazeck FJ, Rescheneder P, von Haeseler A. 2013. NextGenMap: fast and accurate read mapping in highly polymorphic genomes. *Bioinformatics* **29**:2790–2791. doi:10.1093/bioinformatics/btt468

Shen X-X, Opulente DA, Kominek J, Zhou X, Steenwyk JL, Buh KV, Haase MAB, Wisecaver JH, Wang M, Doering DT, Boudouris JT, Schneider RM, Langdon QK, Ohkuma M, Endoh R, Takashima M, Manabe R-I, Čadež N, Libkind D, Rosa CA, DeVirgilio J, Hulfachor AB, Groenewald M, Kurtzman CP, Hittinger CT, Rokas A. 2018. Tempo and Mode of Genome Evolution in the Budding Yeast Subphylum. *Cell* **0**. doi:10.1016/j.cell.2018.10.023
